# Structural Basis and Mode of Action for Two Broadly Neutralizing Antibodies Against SARS-CoV-2 Emerging Variants of Concern

**DOI:** 10.1101/2021.08.02.454546

**Authors:** Wenwei Li, Yaozong Chen, Jérémie Prévost, Irfan Ullah, Maolin Lu, Shang Yu Gong, Alexandra Tauzin, Romain Gasser, Dani Vézina, Sai Priya Anand, Guillaume Goyette, Debashree Chaterjee, Shilei Ding, William D. Tolbert, Michael W. Grunst, Yuxia Bo, Shijian Zhang, Jonathan Richard, Fei Zhou, Rick K. Huang, Lothar Esser, Allison Zeher, Marceline Côté, Priti Kumar, Joseph Sodroski, Di Xia, Pradeep D. Uchil, Marzena Pazgier, Andrés Finzi, Walther Mothes

## Abstract

Emerging variants of concern for the severe acute respiratory syndrome coronavirus 2 (SARS-CoV-2) can transmit more efficiently and partially evade protective immune responses, thus necessitating continued refinement of antibody therapies and immunogen design. Here we elucidate the structural basis and mode of action for two potent SARS-CoV-2 Spike (S) neutralizing monoclonal antibodies CV3-1 and CV3-25 that remained effective against emerging variants of concern in vitro and in vivo. CV3-1 bound to the (485-GFN-487) loop within the receptor-binding domain (RBD) in the “RBD-up” position and triggered potent shedding of the S1 subunit. In contrast, CV3-25 inhibited membrane fusion by binding to an epitope in the stem helix region of the S2 subunit that is highly conserved among β-coronaviruses. Thus, vaccine immunogen designs that incorporate the conserved regions in RBD and stem helix region are candidates to elicit pan-coronavirus protective immune responses.

## Introduction

Severe acute respiratory syndrome coronavirus-2 (SARS-CoV-2) is the third coronavirus to enter the human population since 2003 and is responsible for the coronavirus disease of 2019 (COVID-19) pandemic (Dong et al., 2020; Zhu et al., 2020). While over ∼1 billion vaccines have been administered as of today (Baden et al., 2020; Folegatti et al., 2020; Logunov et al., 2021; Polack et al., 2020; Sadoff et al., 2021a; Sadoff et al., 2021b; Voysey et al., 2021), the pandemic remains uncontrolled in many countries and new variants, including the B.1.1.7 (SARS-CoV-2 α), B.1.351 (β), P.1 (γ), and B.1.617.2 (δ), are outcompeting previous variants due to higher transmissibility and elevated immune evasion (Campbell et al., 2021; Hoffmann et al., 2021; Planas et al., 2021a; Planas et al., 2021b; Prévost and Finzi, 2021; Volz et al., 2021). The spike protein (S) on the surface of the virus mediates entry into cells and is a prominent target for the host immune response including neutralizing antibodies. Consequently, S is a main immunogen for vaccine design. The Moderna, Pfizer-BioNTech, Johnson & Johnson and AstraZeneca vaccines are all based on S immunogens (Baden et al., 2020; Folegatti et al., 2020; Polack et al., 2020; Sadoff et al., 2021a; Sadoff et al., 2021b; Voysey et al., 2021). S consists of a trimer of S1/S2 heterodimers. S1 contains the receptor-binding domain (RBD) that interacts with the cellular receptor angiotensin-converting enzyme 2 (ACE2) (Hoffmann et al., 2020; Li et al., 2003; Walls et al., 2020). S2 possesses the fusion machinery, which can mediate host-viral membrane fusion after S1 shedding. Structural insights into the S protein have been gained by single particle cryo electron microscopy (SP cryoEM) of a soluble trimer comprising most of the ectodomain (Walls et al., 2020; Wrapp et al., 2020), as well as by cryo-electron tomography (cryoET) and SP cryoEM of native virus particles (Ke et al., 2020; Turoňová et al., 2020; Yao et al., 2020). These studies have revealed several distinct prefusion conformations wherein three RBD adopt up or down orientations. Receptor ACE2 binds and stabilizes RBD in the up conformation (Lan et al., 2020; Shang et al., 2020; Xiao et al., 2021; Xu et al., 2021). Single molecule fluorescence resonance energy transfer (smFRET) imaging of single spike molecules on the surface of virus particles has provided real-time information for transitions between both RBD-up and down conformations through one necessary intermediate (Lu et al., 2020).

Antibodies isolated from convalescent patients, vaccinated individuals and previous work on the related SARS-CoV-1 and MERS-CoV viruses can be classified by their specificity for three main epitopes: the RBD, the N-terminal domain (NTD) and the S2 subunit (Barnes et al., 2020b; Jennewein et al., 2021; Ju et al., 2020; Liu et al., 2020; Montefiori and Acharya, 2021; Ullah et al., 2021). For each class the conformational preferences for either RBD-up or RBD-down trimer configurations have been described. Antibodies directed against the RBD are often attenuated against emerging variants of concern due to escape mutations (Greaney et al., 2021a; Liu et al., 2021; Starr et al., 2021; Weisblum et al., 2020). Although immune responses elicited by existing vaccines do offer protection to varying degrees against all known variants of concern (Skowronski et al., 2021; Tauzin et al., 2021), a booster shot to ensure sufficient protection from future emerging variants might be needed. Moreover, SARS-CoV-2 is the third β-coronavirus after SARS-CoV-1 and MERS-CoV to be transferred to humans in the 21^st^ century and given the large natural reservoir of similar viruses in species such as bats (Anthony et al.; Ge et al., 2013; Letko et al., 2020; Menachery et al., 2015; Menachery et al., 2016; Wang et al., 2018), another pandemic caused by a new coronavirus is likely to happen again. These coronaviruses possess a conserved S2 domain, which raises the possibility of cross-reactive antibodies and cross-reactive vaccines. SARS-CoV-2 S is approximately 75% homologous to SARS-CoV-1 and 35% to MERS S (Grifoni et al., 2020; Zhou et al., 2020). Various cross-reactive antibodies have been identified (Hoffmann et al., 2020; Jennewein et al., 2021; Ma et al., 2020; Ng et al., 2020; Song et al., 2020; Tian et al., 2020; Wang et al., 2020; Wang et al., 2021). Recently isolated antibodies capable of cross-neutralizing human coronaviruses bind to the conserved stem helix region on S2, reviving hopes for pan-coronavirus vaccines (Pinto et al., 2021; Sauer et al., 2021; Zhou et al., 2021).

We previously characterized two potent S-binding antibodies, CV3-1 and CV3-25, out of 198 antibodies isolated from convalescent patients (Jennewein et al., 2021; Ullah et al., 2021). CV3-1 targets the RBD of S1 and CV3-25 binds to the S2 ectodomain, the former displaying the most potent neutralizing activity among all antibodies (Abs) isolated. While CV3-1 is specific for the RBD of SARS-CoV-2, CV3-25 can recognize the S2 domains derived from several β-coronaviruses (Jennewein et al., 2021). Both antibodies protected against SARS-CoV-2 in animal models in prophylactic and therapeutic settings (Ullah et al., 2021). Here we report the structural basis and mode of action for these two potent antibodies. We deployed cryoET of virus like particles (VLPs) carrying the S_B.1.1.7_ variant to determine the epitopes of these two antibodies. CV3-1 bound to the tip region (485-GFN-487 loop) within the receptor-binding motif (RBM), as confirmed by mutagenesis. Interestingly, we observed that most spikes in CV3-1-treated virus-like particles (VLP) were triggered into the post-fusion conformation of S2 and caused S1 shedding into the supernatant. The data indicate that CV3-1 is a potent agonist and point to the 485-GFN-487 loop as an allosteric center critical for the activation of S1. In contrast, CV3-25 bound the stem helix in the connecting domain (CD) of S2 and blocked membrane fusion. Its binding was asymmetric as S trimer was bound by 1 or 2 CV3-25 antigen-binding fragments (Fabs). Peptide competition narrowed the epitope and permitted the determination of the crystal structure of the S2 stem peptide bound to CV3-25 Fab. The structure revealed a unique bent conformation of the viral peptide with an upstream α-helical region followed by a random coil. Fitting of the X-ray structure into the cryoET density map demonstrated that an increasing degree of stem helix rotation was required to allow binding of one or both Fabs to avoid steric clashes. Compared to other recently reported stem helix engaging antibodies, the advantage of CV3-25 appears to be that it binds to the flexible loop, likely facilitating stem helix engagement. Given that the stem helix epitope is highly conserved among β-coronaviruses, immunogens featuring this S2 epitope are interesting candidates for vaccines to cover all variants and possibly exhibit pan-coronavirus efficacy. Moreover, since many antibodies that bind spike are non-neutralizing, our work suggests that agonist features that prematurely trigger and thereby irreversibly inactivate S, or inhibition of membrane fusion, contribute to the ability of neutralizing antibodies to block SARS-CoV-2 infection.

## Results

### CV3-1 and CV3-25 Neutralize Emerging SARS-CoV-2 Variants

We first tested the ability of CV3-1 and CV3-25 to recognize and neutralize the emerging variants of concern, B.1.1.7 (SARS-CoV-2 α), B.1.351 (β), P.1 (γ), B.1.617.2 (δ) as well as variants of interest B.1.429 (ε), B.1.525 (η), B.1.526 (ι) and B.1.617.1 (κ). CV3-1 efficiently bound to cells expressing spike proteins from these different SARS-CoV-2 variants or carrying their individual mutations (Figure 1A & S1A). Despite the presence of variant-specific mutations in RBD, CV3-1 retained potent neutralizing activity (IC_50_ 0.004-0.014 μg/ml) (Figure 1B). Of note, CV3-1 binding to the B.1.1.7 variant with and without the additional E484K substitution was higher than binding to the spike from the original Wuhan strain (WT). CV3-25 was less potent with an IC_50_ in the range of ∼0.05-0.2 μg/ml, but remained effective against all variants in both binding ability and neutralization (Figure 1A, B, Figure S1B). Both CV3-1 IgG and the CV3-25 IgG GASDALIE mutant, which binds more strongly to Fcγ receptors, also protected *in vivo* against both the B.1.1.7 (α) (Ullah et al., 2021) and B.1.351 (β) variants of SARS-CoV-2 in the K18-hACE2 prophylactic mouse model (Figure 1C-E). Both antibodies limited viral replication in the nose and lungs as well as its dissemination to the brain, thereby reducing the induction of pro-inflammatory cytokines (Figure S1C-F). These data demonstrate that in contrast to other antibodies that are attenuated against emerging variants (Greaney et al., 2021a; Liu et al., 2021; Starr et al., 2021; Weisblum et al., 2020), CV3-1 and CV3-25 remain potent against these variants and are therefore prime candidates to elucidate the mode of action and identify epitopes with pan-coronavirus activity.

**Figure 1.**
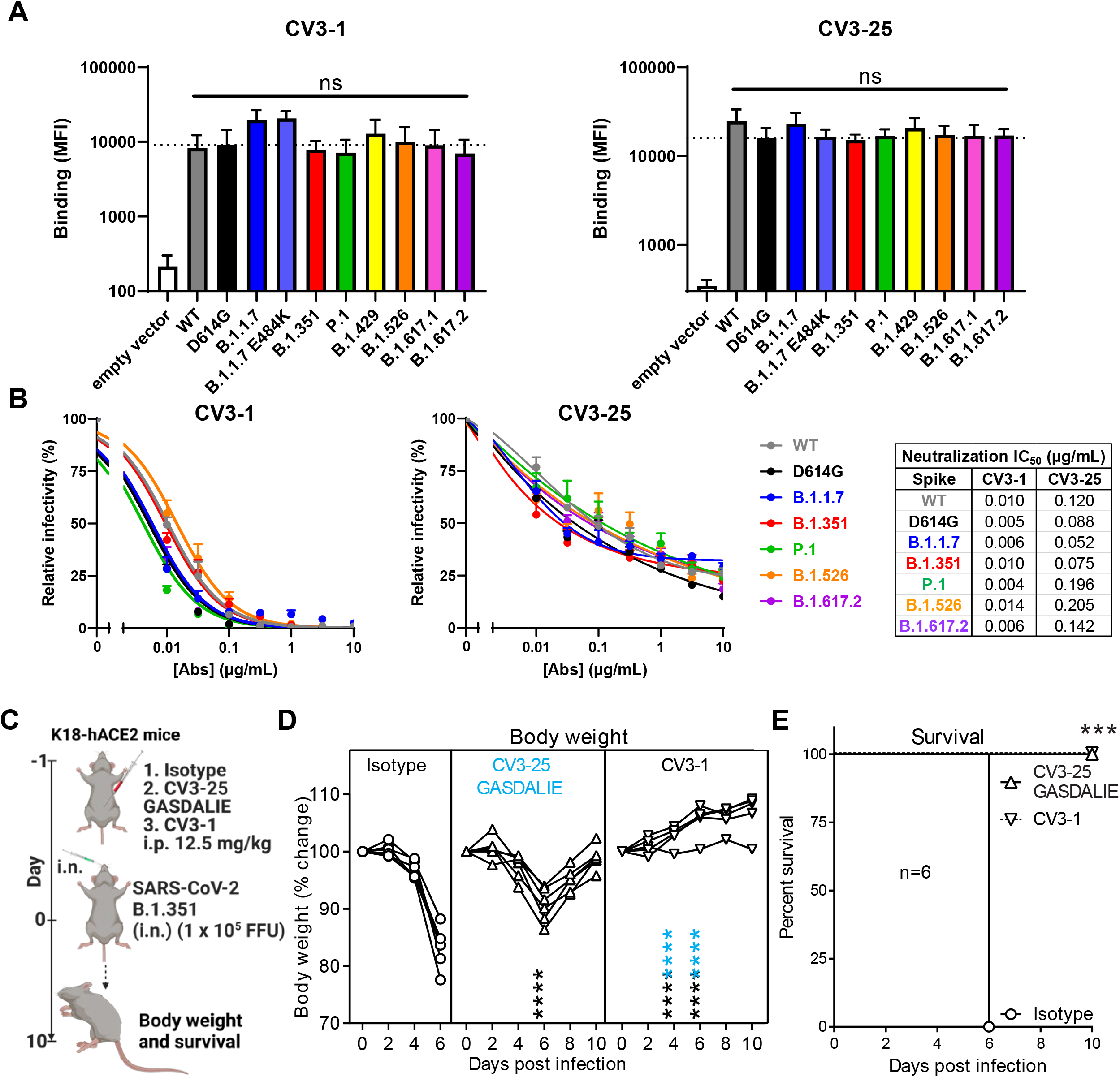
CV3-1 and CV3-25 Neutralize Emerging SARS-CoV-2 Variants. (**A**) Cell-surface staining of 293T cells expressing full-length Spike from indicated variants by CV3-1 (left panel) and CV3-25 (right panel) monoclonal Abs (mAbs). The graphs show the median fluorescence intensities (MFIs). Dashed lines indicate the reference value obtained with Spike D614G. Error bars indicate means ± SEM. These results were obtained in at least 3 independent experiments. Statistical significance was tested using one-way ANOVA with a Holm-Sidak post-test (ns, non significant). (**B**) The ability of CV3-1 and CV3-25 mAbs to neutralize Wuhan-Hu-1 (WT), D614G mutant, B.1.1.7, B.1.351, P.1, B.1.526 and B.1.617.2 pseudoviruses infectivity in 293T-hACE2 cells was measured as indicated in Star Methods. IC_50_ values are shown. Error bars indicate means ± SEM. These results were obtained in at least 3 independent experiments. (**C**) A scheme showing the experimental design for testing the in vivo efficacy of NAbs, CV3-1 WT and CV3-25 G236A/S239D/A330L/I332E (GASDALIE) mutant (12.5 mg IgG/kg body weight) delivered intraperitoneally (i.p.) 1 day before challenging K18-hACE2 mice with a lethal dose (1 x 10^5^ FFU) of B.1.351 SARS-CoV-2. Human IgG1-treated (12.5 mg IgG/kg body weight) mice were used as control. (**D**) Temporal changes in mouse body weight in experiment shown in (C), with initial body weight set to 100%. (**E**) Kaplan-Meier survival curves of mice (n = 6 per group) statistically compared by log-rank (Mantel-Cox) test for experiments as in (C). Grouped data in (D) were analyzed by 2-way ANOVA followed by Tukey’s multiple comparison tests. Statistical significance for group comparisons to isotype control are shown in black and for those to CV3-25 GASDALIE are shown in blue., p < 0.05;, p < 0.01;, p < 0.001;, p < 0.0001; Mean values ± SD are depicted.

### Spike Conformational Preferences of CV3-1 and CV3-25 Assessed by smFRET and CryoET

We utilized smFRET as a dynamic method and cryoET as a static method to characterize the conformational preferences of CV3-1 and CV3-25 for S of the B.1.1.7 variant (S_B.1.1.7_). smFRET measures the conformational state within a single S1 protomer and indicated that the unliganded S_B.1.1.7_ has access to 4 distinct conformational states, with the ∼0.5 FRET state being the most occupied state (Figure 2A). We had previously established that these states correspond to the RBD-down (∼0.5 FRET), RBD-up (∼0.1 FRET), a necessary structural intermediate (∼0.3 FRET) in the transition from RBD-down to RBD-up that is likely observed in a protomer adjacent to an RBD-up, and a high-FRET state (∼0.8) for which a structure is not available (Lu et al., 2020). CV3-1 redistributed the conformational landscape of S to the ∼0.1 low FRET state that corresponds to the RBD-up thus mimicking receptor ACE2. CV3-25 redistributed the conformational landscape towards activation with an increase in the occupancy of the structural intermediate (∼0.3 FRET) as well as the RBD-up state (∼0.1 FRET) (Figure 2A, B). Overall, the conformational landscapes of the S_B.1.1.7_ variant and the conformational preferences of CV3-1 and CV3-25 were similar to the original Wuhan strain (Ullah et al., 2021).

**Figure 2.**
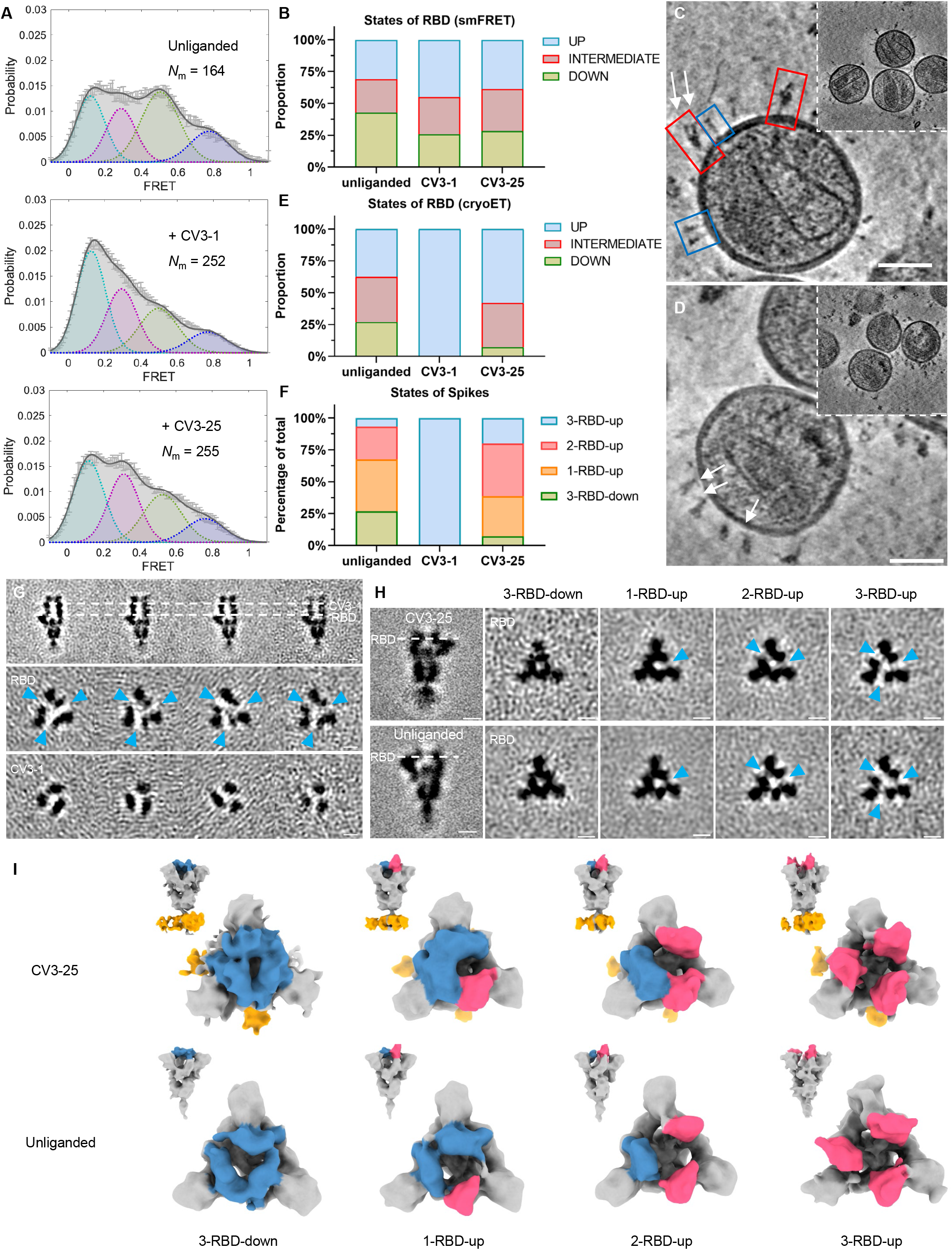
Conformational Dynamics of S_B.1.1.7_ Bound with CV3-1 and CV3-25. (**A**) Conformational states of S_B.1.1.7_ on lentiviral particles monitored by smFRET for unliganded, CV3-1 and CV3-25 bound S_B.1.1.7_. FRET histograms with number (N_m_) of individual dynamic molecules/traces compiled into a conformation-population FRET histogram (gray lines) and fitted into a 4-state Gaussian distribution (solid black) centered at 0.1-FRET (dashed cyan), 0.3-FRET (dashed red), 0.5-FRET (dashed green), and 0.8-FRET (dashed magenta). (**B**) Proportion of different states of RBD identified by smFRET in (A). For parallel comparation to cryoET data, 0.8-FRET portion was omitted, due to its structural uncertainty. (**C**-**D**) Zoomed-in views of SARS-CoV-2 pseudoviruses bearing S bound by CV3-1 (C) and CV3-25 (D) Fabs and representative slices of tomograms (insets). Scale bar, 50 nm. White arrows indicate bound Fabs. Red boxes, prefusion spikes; Blue boxes, post-fusion spikes. (**E**) Proportion of different states of RBD at different conditions from cyroET data. UP state was separated by focused classification on the RBD region. The remaining of the RBDs were defined as DOWN state if there was no up RBD on the same spike, otherwise, they were considered as INTERMIEDATE state. (**F**) Proportion of different RBD states of spikes on virions with and without Fabs bound. Spikes were grouped into 3-RBD-down,1-RBD-up, 2-RBD-up and 3-RBD-up classes. (**G**) Side views (top panel) and top views (middle and bottom panels) of subclasses of averaged S bound by CV3-1 Fabs. (**H**) Side views (left column) of the consensus structure of unliganded (bottom) and CV3-25 bound (top) S and top views of subclass averages (right columns) obtained after focused classification on the RBD of S. In (G-H), dotted lines indicate the positions of top-view sections. Blue arrowheads point to the gap in density between RBD and the neighboring NTD that appears when the RBD moves into the UP-state. Scale bar, 5 nm. (**I**) Segmentation of subclass averages of unliganded (bottom) and CV3-25 bound (top) S. Top views and side views (insets) are shown for 3-RBD-down, 1-RBD-up, 2-RBD-up and 3-RBD-up classes. Down RBDs and up RBDs are shown in blue and red respectively, CV3-25 Fabs are shown in orange.

We next used cryoET to identify the epitope for CV3-1 and CV3-25 and analyze their conformational preference by quantifying the proportion of antibody-bound trimers in the 3-RBD-down, 1-RBD-up, 2-RBD-up and 3-RBD-up for S_B.1.1.7_ on the surface of lentiviral particles. To improve incorporation of S into lentiviral particles for EM, the S_B.1.1.7_ cytoplasmic tail was truncated (Figure 2C, D). The unliganded S_B.1.1.7_ displayed a similar number of 3-RBD-down, 1-RBD-up, and 2-RBD-up conformations, with the 3-RBD-up conformation rarely observed (Figure 2F). CV3-1 clearly bound to the top of RBD with the RBD being oriented up (Figure 2C). Nearly all trimers with bound CV3-1 were in the RBD-up conformation (Figure 2E-G). Binding to RBD is consistent with previous data that demonstrated the ability of CV3-1 to competitively inhibit ACE2-S binding in vitro (Jennewein et al., 2021).

In contrast, CV3-25 bound towards the bottom of S2 and all trimer configurations were observed (Figure 2D, F, H, I). Compared to the unliganded S, CV3-25 binding redistributed the frequency of trimer configurations from the 3-RBD-down to the 1-, 2-, and 3-RBD-up configurations. To compare cryoET to smFRET data, we calculated the number of conformational states of individual RBD units, which is monitored by smFRET. This was done under the assumption that protomers neighboring to an RBD-up protomer are in an intermediate FRET state (Lu et al., 2020). Consequently, the 1-& 2-RBD-up not only feature 1 or 2 additional protomers in the RBD-up conformation, but also likely introduce a significant occupancy for structures exhibiting an intermediate FRET state (∼0.3) (Figure 2E, F). While several caveats remain, such as the use of cytoplasmic tail-deleted S for EM (wt S for smFRET), the inability to see a structure corresponding to the intermediate FRET (∼0.3), and as a consequence not knowing if both, left and right protomers neighboring a RBD-up, are in an intermediate FRET state, and the inability to assign a structure for the high-FRET state (∼0.8), this is the first time that we can generate dynamic and static data for S on virus particles produced in the same cell type and assess them in parallel by smFRET and cryoET. Overall, there is qualitative agreement between cryoET and smFRET about how CV3-1 and CV3-25 alter the conformational landscape of S. Above mentioned caveats make quantitative comparisons currently impossible. smFRET may detect more dynamic features while cryoET may emphasize static features as previously discussed for the HIV-1 spike protein (Li et al., 2020).

### CV3-1 Binds to the 485-GFN-487 Loop of RBD

To gain a higher resolution structure for CV3-1 bound to S_B.1.1.7_ we imposed C3 symmetry on a subtomogram averaged structure and determined a ∼12 Å map (Figure 3A, B, Figure S3A-C). The averaged cryoET structure showed three CV3-1 Fabs bound to the apex of the S trimer. Classification among these particles did not identify any subclass of spikes bound with only one or two CV3-1 Fabs (Figure 2D). Rigid-body fitting with a 3-RBD-down atomic model of S_B.1.1.7_ (PDB: 7LWS (Gobeil et al., 2021)) left all three RBDs outside of cryoET density, while flexible fitting resulted in the conformational change from the RBD-down to the RBD-up state (Figure 3C, Video S1). We applied rigid fitting of the atomic structure of 1-up RBD (PDB: 7LWV (Gobeil et al., 2021)) to arrive at a model for CV3-1 Fab bound to S_B.1.1.7_ (Figure 3D). Compared to the footprint of receptor ACE2 on RBD (PDB: 7KJ4 (Xiao et al., 2021)), CV3-1 preferentially bound to the extending loop that contains the G^485^F^486^N^487^ residues (Figure 3D-E). We performed mutagenesis for the RBM and tested the abilities of CV3-1 and ACE2 to bind S mutants expressed on cells by flow cytometry. In agreement with the structural model, CV3-1 binding was preferentially affected by mutations in the 485-GFN-487 loop (Figure 3F, G). In contrast, ACE2 binding was sensitive to mutations within the RBM consistent with previous results (Greaney et al., 2021b; Starr et al., 2020). Importantly, all mutations within the 485-GFN-487 loop affecting CV3-1 binding, also impaired ACE2 binding indicating that escape mutations at these positions would likely result in a high fitness cost for the virus.

**Figure 3.**
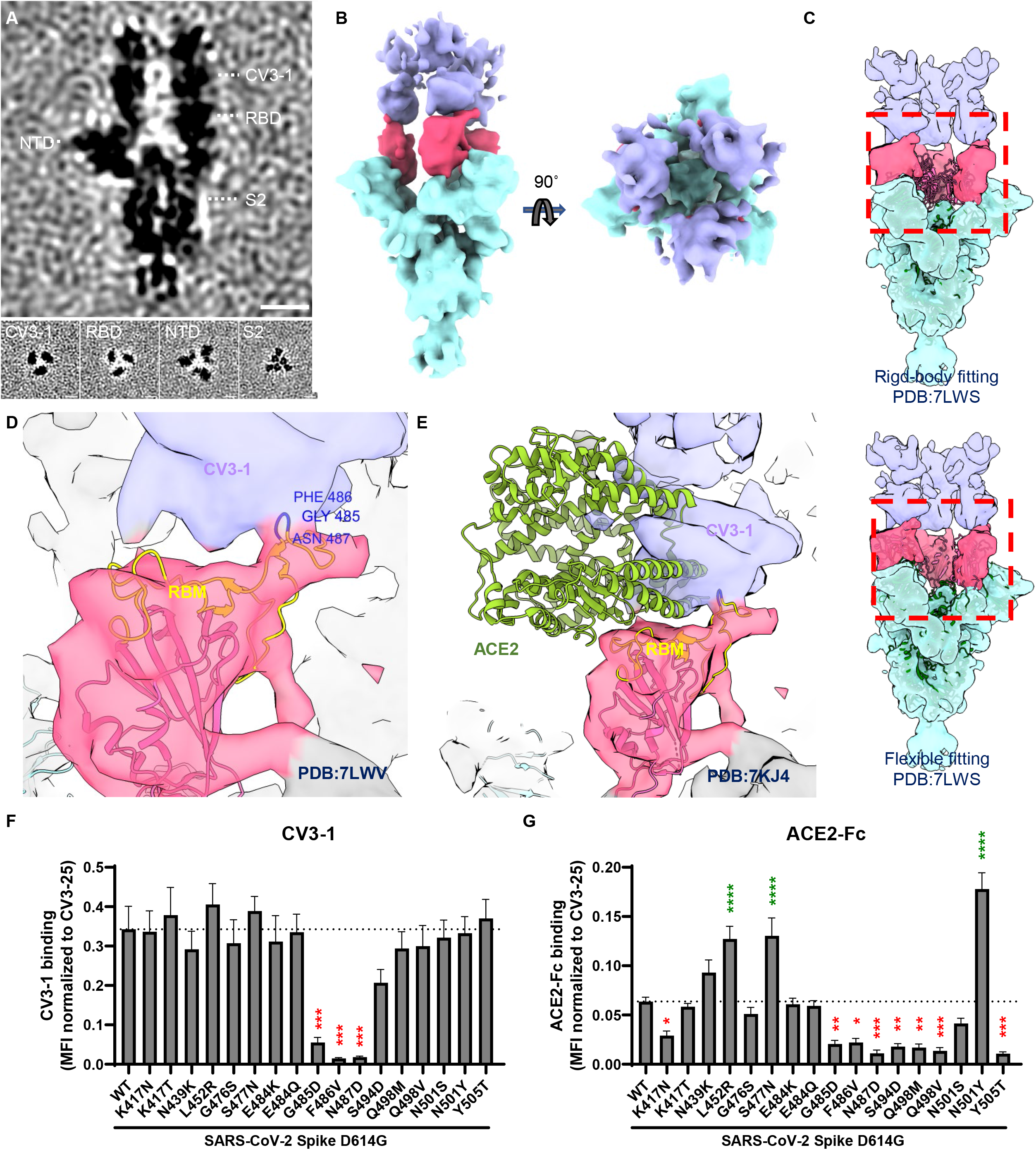
CV3-1 Binds to the 485-GFN-487 Loop of RBD. (**A**) Side view (top panel) and top views (bottom panel) of subtomogram average of CV3-1 bound S. Dotted lines indicate the positions of top-view sections. Scale bar, 5 nm. (**B**) Segmentation of CV3-1 bound S. Side view (left) and top view (right) are shown. CV3-1 Fabs are shown in purple, RBDs are shown in red and the rest of S in cyan. (**C**) Fitting cryoET density map of CV3-1 bound S with 3-RBD-down atomic model of S (PDB: 7LWS). Top panel: rigid-body fitting. Bottom panel: flexible fitting. (**D, E**) Zoomed-in view of cryoET map fitting with RBD-up atomic model (D, PDB: 7LWV) and ACE2-S atomic model (E, PDB: 7KJ4) at the interaction site. (**F-G**) Binding of CV3-1 (F) and ACE2-Fc (G) to 293T cells expressing selected full-length Spike harboring RBM mutations. The graphs shown represent the median fluorescence intensities (MFIs) normalized to the MFI obtained with CV3-25 staining of the corresponding mutant. Dashed lines indicate the reference value obtained with Spike D614G (WT). Error bars indicate means ± SEM. These results were obtained in at least 4 independent experiments. Statistical significance was tested using one-way ANOVA with a Holm-Sidak post-test (*p < 0.05; **p < 0.01; ***p < 0.001; ****p < 0.0001).

### CV3-1 is a Potent Agonist Triggering S1 Shedding

SARS-CoV-2 S proteins lacking the cytoplasmic tail are efficiently incorporated into lentiviral particles and form a dense array of spikes in the prefusion state (Dieterle et al., 2020; Ou et al., 2020; Schmidt et al., 2020; Yu et al.). In contrast, the virus particles incubated with CV3-1 lost most prefusion spikes and displayed S in the post-fusion state (Figure 2C). Quantification of spike numbers revealed that 83% of prefusion spikes, comparing to unliganded S, were lost after incubation with CV3-1 (Figure 4A). The structural characterization of CV3-1 bound to S shown above was performed with the remaining ∼17% of prefusion spikes. Given the loss of S1 and activation of S2 into post-fusion conformation, we hypothesized that, besides competition with ACE2 (Jennewein et al., 2021), triggering S1 shedding likely contributed to SARS-CoV-2 neutralization efficacy of CV3-1. Radioactive labeling followed by immunoprecipitation of cell lysates and supernatant revealed that incubation with CV3-1 indeed released most S1 into the supernatant with its activity well exceeding that of ACE2 (Figure 4B). S lacking the furin-cleavage site was resistant to CV3-1-and ACE2-mediated shedding. The loss of S1 following incubation of CV3-1 was also observed by flow cytometry on cells expressing S (Figure 4C). S lacking the furin-cleavage site was again resistant to shedding induced by CV3-1. In contrast to CV3-1, CV3-25 induced little or no shedding in all assays (Figure 4A-D). Importantly, the ability of CV3-1 to neutralize the emerging variants B.1.1.7 (SARS-CoV-2 α), B.1.351 (β), P.1 (γ), B.1.526 (ι), B.1.429 (ε) and B.1.617.2 (δ) (Figure 1B) paralleled the ability of CV3-1 to shed S1 (Figure 4D). These data indicate that RBD-targeting antibodies can be potent agonists by prematurely activating S to impair virus entry.

**Figure 4.**
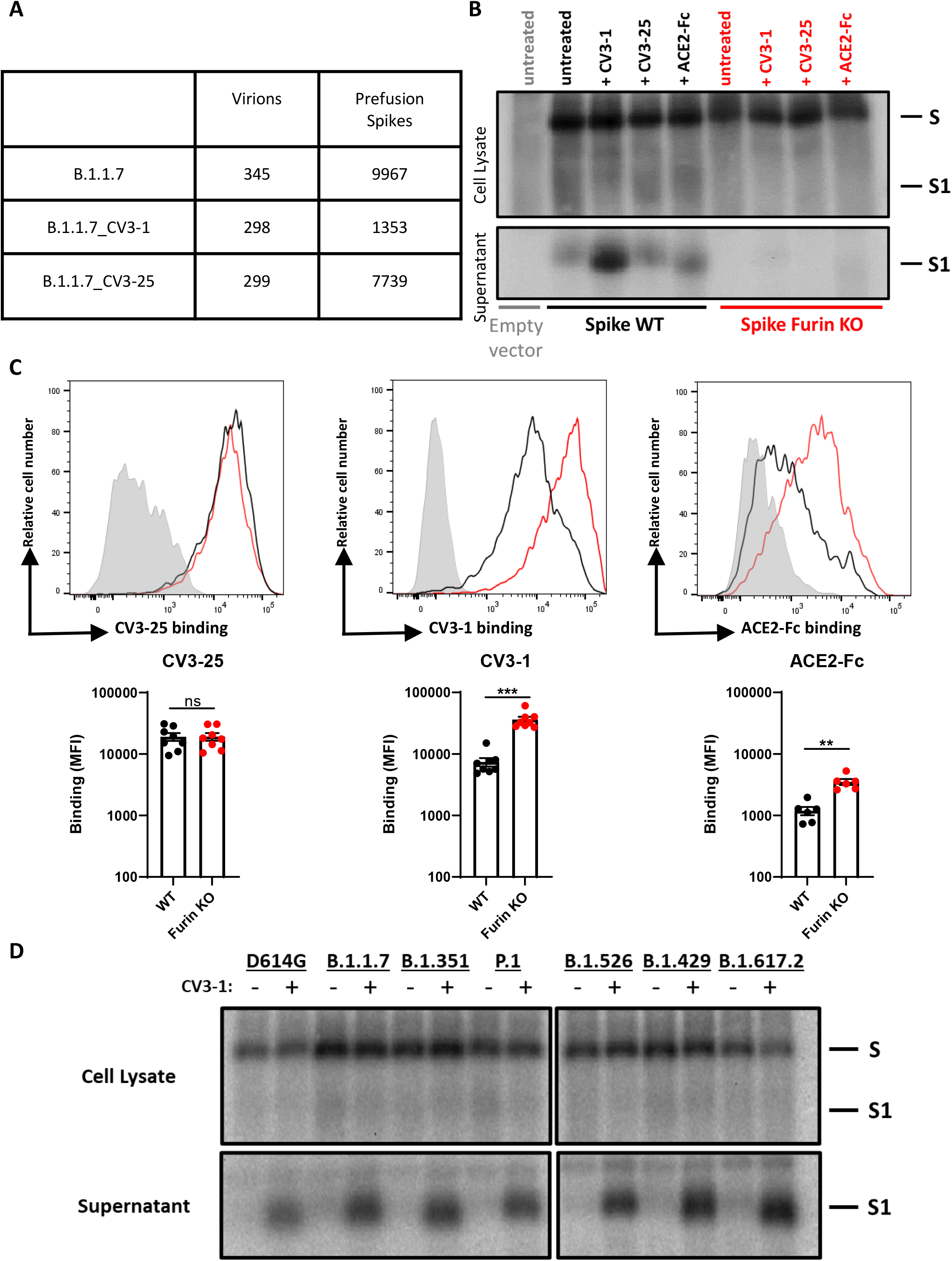
CV3-1 Triggers S1 Shedding. (**A**) Statistical table of pre-fusion S selected manually for cryoET analysis. (**B**) S1 shedding was evaluated by transfection of 293T cells followed by radiolabeling in presence of CV3-1, CV3-25 or ACE2-Fc and immunoprecipitation of cell lysates and supernatant with CV3-25 and a rabbit antiserum raised against SARS-COV-2 RBD produced in-house. Furin KO = furin cleavage site knockout. These results are representative of two independent experiments. (**C**) CV3-25, CV3-1 and ACE2-Fc recognition of 293T cells expressing the full-length SARS-CoV-2 ancestral S with or without (Furin KO) a functional furin cleavage site. Histograms depicting representative cell-surface staining of cells transfected with wild-type Spike (black line), Furin KO (red line) or with an empty vector (light gray). Error bars indicate means ± SEM. These results were obtained in at least 6 independent experiments. Statistical significance was tested using a Mann-Whitney U test (**p < 0.01; ***p < 0.001; ns, non significant). (**D**) CV3-1 induced S1 shedding of S from selected emerging variants, measured as in (B).

As previously shown using SP cryoEM, the S trimer displays significant tilt relatively to the viral membrane because of its highly flexible stalk region (Ke et al., 2020; Turoňová et al., 2020; Yao et al., 2020). Among the CV3-1-bound S that remained on the surface of virus particles, we sought to observe a change in the tilt angle of CV3-1-bound S. Quantification revealed a profound straightening of the S from an average tilt angle of ∼57° for the unliganded S to only ∼37° (Figure S2A-C). Apparently, the ACE2–mimicking activation of RBD by CV3-1 leads to long-range structural effects involving S2, likely weakening the S1-S2 interface and resulting in the shedding of S1. When evaluating antibody binding cooperativity, the Hill coefficient for CV3-1 binding was found to be highly positive (h>2) (Figure S2D). This highly positive cooperativity may suggest that CV3-1 could access its epitope in the down conformation and bring the RBD to the up conformation to facilitate exposure of neighboring subunits, consistent with previous observations seen with other class 2 RBD Abs (Barnes et al., 2020a; Brouwer et al., 2020).

### CV3-25 Binds the Stem Helix of S2

We employed a multipronged approach including cryoEM, cryoET, peptide competition, and X-ray crystallography to gain mechanistic insight into how CV3-25 achieves broad neutralization against emerging SARS-CoV-2 variants and other β-coronaviruses (Jennewein et al., 2021; Ullah et al., 2021). We first determined the cryo-EM structure of the SARS-CoV-2 spike (HexaPro, prefusion-stabilized)(Hsieh et al., 2020) in the presence of CV3-25 Fab at an overall resolution of ∼3.5 Å (Figure S4A-D). Map density analysis indicated the 1-RBD-up state was the dominant spike conformation with a decreased local resolution in this region (Figure S4G). The density corresponding to the C-terminal stem region was less defined with a local resolution lower than 7 Å, but there was additional density for CV3-25 in the C-terminal stem region. 3D classification of the Cryo-EM data barely improved the local density suggesting incomplete Fab saturation for all available binding sites. Nevertheless, the data suggested that CV3-25 binds to the lower stem of the soluble HexaPro S.

Given that soluble S trimers are truncated, lack the transmembrane region, and feature a T4 foldon, we reasoned that cryoET of native spike proteins embedded into virus particles could provide more insight into CV3-25’s epitope. We used cryoET followed by subtomogram averaging of ∼7000 prefusion spikes to examine CV3-25 binding to S_B.1.1.7_. Subclassification revealed that about half of S had two CV3-25 Fabs bound to the stem of S2, and the other half had only one CV3-25 Fab bound (Figure S5). We further aligned the subtomograms with a mask for two CV3-25 Fabs to arrive at ∼10 Å resolution map. This structure places the CV3-25 epitope within the connecting domain (CD) of the stem helix (Figure 5A, B, F). Density for the second Fab was weaker since ∼half of spikes had only one CV3-25 Fab bound.

**Figure 5.**
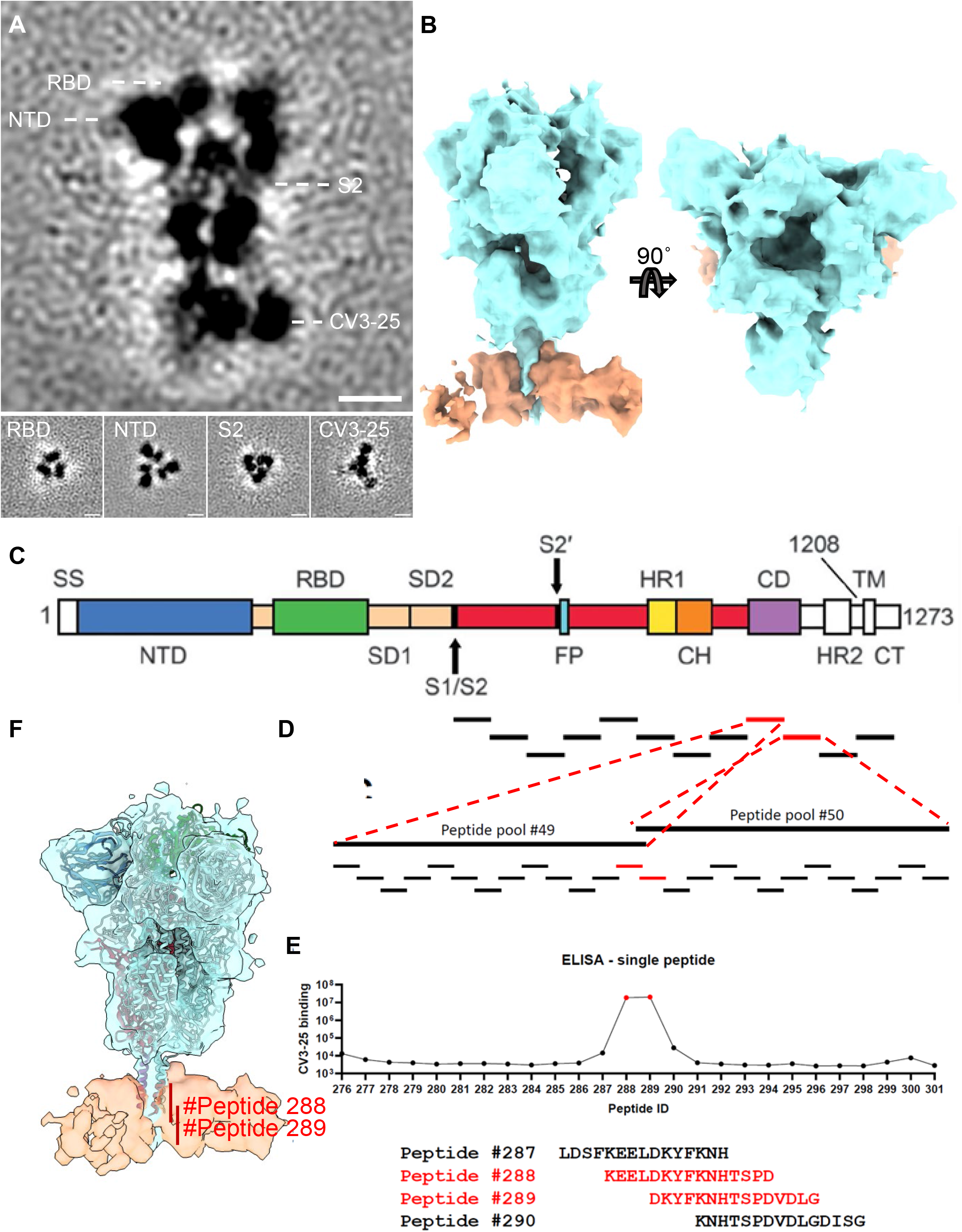
CV3-25 Binds to a Conserved Epitope in S2. **(A)** Side view (top panel) and top views (bottom panel) of subtomogram averaged CV3-25 bound S. Dotted lines indicate the positions of top-view sections. Scale bar, 5 nm. **(B)** Segmentation of CV3-25 bound S. Side view (left) and top view (right) are shown. CV3-25 Fabs are shown in orange and S is shown in cyan. **(C)** SARS-CoV-2 Spike sequence depicting the different subunits and domains composing the full-length Spike protein. With permission from AAAS. (Wrapp et al., 2020) **(D)** Pools of peptide covering the whole S2 subunit sequence were used to identify the linear region recognized by CV3-25 mAb. Indirect ELISA was performed using SARS-CoV-2 S2 peptide pools and incubation with the CV3-25 mAb. Peptide pools covering the connector domain (CD) region with significant positive signal were highlighted in red (peptide pools #49 and #50). Depiction of the SARS-CoV-2 Spike individual peptides from the peptide pools #49 and #50, with a 4 amino acid residue overhang. Individual peptides covering the S2 connector domain region were used to identify the region recognized by CV3-25 mAb. **(E)** Indirect ELISA was performed using SARS-CoV-2 S2 individual peptides (from peptide pools #49 and #50) and incubation with the CV3-25 mAb. CV3-25 binding was detected using HRP-conjugated anti-human IgG and was quantified by relative light units (RLU). Single peptides with significant positive signal were highlighted in red (peptides #288 and #289). Amino acid sequence of peptides recognized by CV3-25 (peptides #288 and #289, shown in red) and of neighboring peptides not recognized by CV3-25 (peptides #287 and #290, shown in black). **(F)** Rigid fitting cryoET density map of CV3-25 bound S with atomic model of closed prefusion S (PDB:6XR8). Peptides #288 and #289 location at the CV3-25 binding site are indicated in red.

As discussed above, classification of the spike structures into 3-RBD-down, 1-, 2-or 3-RBD-up revealed an overall shift towards activation for CV3-25 bound S (Figure 2E, F). Averaged structures focusing on RBD showed 3 bound CV3-25 Fabs in 1-RBD-up and 2-RBD-up spikes while only 2 CV3-25 Fabs bound to 3-RBD-down and 3-RBD-up average structures (Figure S2E). These structures confirmed that CV3-25 binds to all prefusion spike configurations consistent with previous biochemical results as well as smFRET (Lu et al., 2020). Any observed asymmetry was not due to the tilt of the spike as the average tilt barely changed upon binding of CV3-25 (Figures S2A-C), consistent with a neutral antibody binding cooperativity (Hill coefficient ≈ 1) (Figure S2D).

### Peptide Screening Maps CV3-25 Epitope to the Spike Residues 1149-1167

To gain atomic insight, we screened S2 peptides for binding to CV3-25 with the goal of isolating peptides suitable for X-ray crystallography. The first insight that CV3-25 binds a linear peptide was gained from Western blotting following SDS-PAGE. CV3-25 was clearly able to bind to S2 as well as the S2-containing S precursor under fully denaturing conditions and independently of N-linked glycans (Figure S6A, B). We then tested a set of peptides (15-mer) spanning the entire S2 subunit including the connecting domain and performed two rounds of ELISA to identify peptides capable of binding CV3-25 (Figure 5C-E). The identified peptides (#288 and #289) were also tested in competition assays and the binding was quantified using SPR assays (Figures S6C, D). Peptide #289 was the most potent in all assays with a K_D_ of 29 nM and efficiently blocked CV3-25 neutralization (Figure S6E, 6F). Peptides #288 and #289 mapped to the S2 stem helix region (Figure 5F), consistent with the CV3-25 binding region indicated in the cryoET averaged structure.

### CV3-25 Binds to a Conserved S2 Peptide in a Bent Conformation

To obtain molecular insight into CV3-25 interaction with the S2 stem peptide, we determined the co-crystal structure of CV3-25 Fab with a synthetic peptide spanning residues 1140-1165 (26mer) of SARS-CoV-2 spike. The structure was solved to 2.1 Å resolution and allowed us to resolve 20 of the 26 residues in relation to the Fab paratope (Figure 6, Figure S7, Table S2). When bound to CV3-25 Fab, the peptide adopted a bent conformation with the N-terminal half of the peptide (residues 1146-DSFKEELDKYFK-1157) forming an α-helix and the C-terminal half a random coil (residues 1158-NHTSPDVD-1165) with a bend of ∼95° between the two (Figure 6A). This bent conformation fit well with the long complementary determining region (CDR) H3 loop of the Fab (16 aa long) that is stabilized by extensive H-bonds, salt bridges and intra-molecular π-π stacking between residues Y^1155^ and H^1159^ of the peptide (Figure 6B-D and Figure S7). A rare CDR H3 disulfide bond between residues C^99^ and C^100D^ also stabilizes the CDR H3 hairpin that tightly associates with the S2 peptide random coil. Interestingly, the S2 stem region recognized by CV3-25 is conserved among the B lineage of β-coronaviruses (Figure 6F), with several key epitope residues also conserved among A, C, and D lineages. This suggests that CV3-25 displays cross-reactivity with coronaviruses beyond the B-lineage. Furthermore, although crystallographic analyses confirm residues 1149 to 1165 of the S2 stem to interact with CV3-25 (Figure 6D), SPR analyses using S2 peptide truncations indicate that CV3-25 may also interact with residues following D^1165^, the terminal S2 residue used in crystallographic studies. These contacts could be mediated by the light chain of CV3-25 that is positioned to accommodate the C-terminal extension of the peptide (Figure 6A). Of note, the S2 recognition site and angle of approach of CV3-25 differentiate it from B6, the only reported anti-MERS-CoV S2 NAb as well as CC40.8 and S2P6, the only two known human anti-SARS-CoV-2 S2 NAbs (PDBs not available) (Pinto et al., 2021; Sauer et al., 2021; Zhou et al., 2021) (Figure 6G). B6, CC40.8 and S2P6 mainly interact with the N-terminal stem α-helix and barely contact with C-terminal loop as recognized and reconfigured by CV3-25, indicating that CV3-25 is the first representative of a new class of anti-S2 antibodies with broad reactivity against β-coronaviruses.

**Figure 6.**
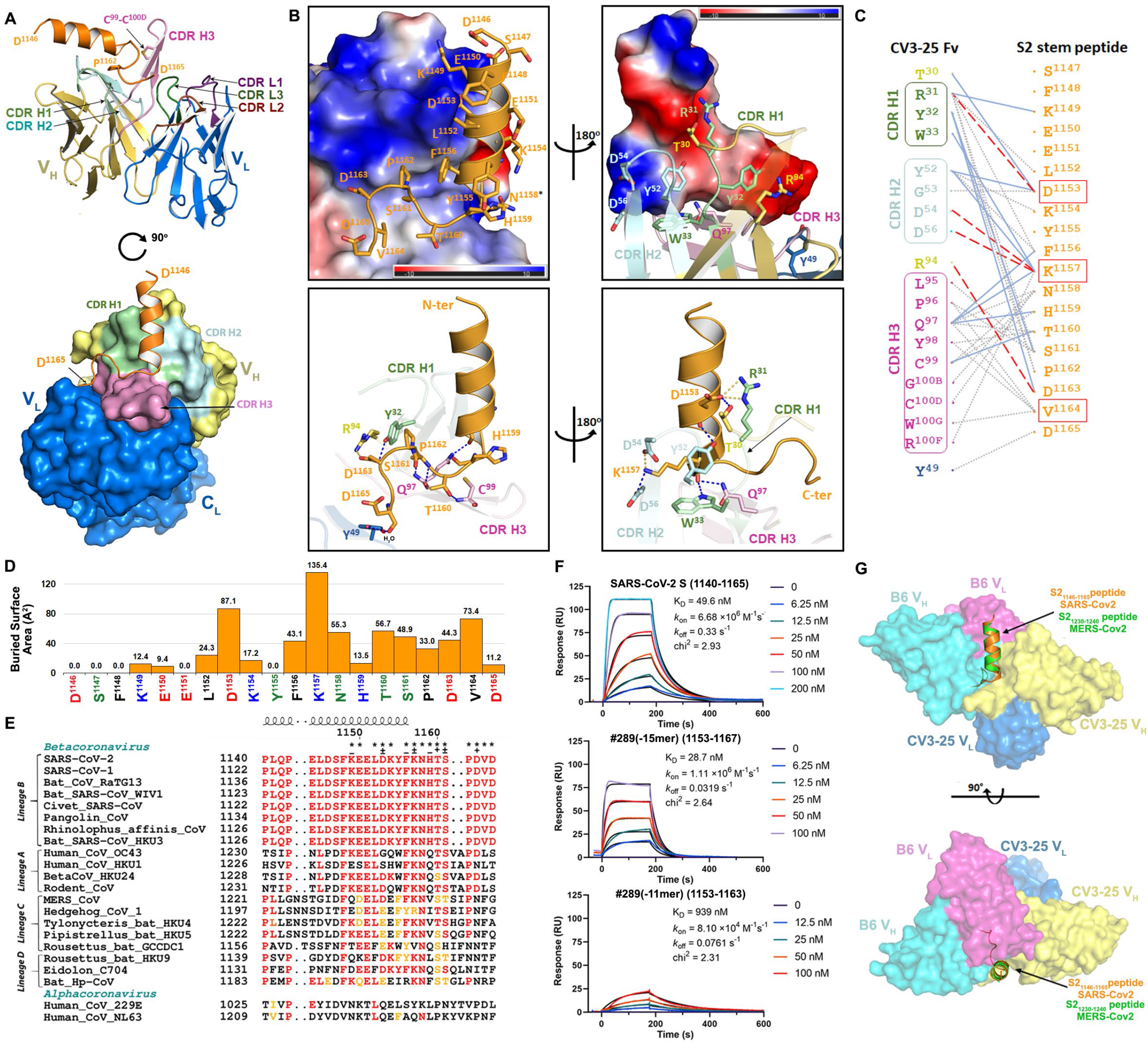
Molecular Details of Interaction of CV3-25 with SARS-CoV-2 Stem Peptide Spanning Residues 1140-1165 of S2. (**A**) Crystal structure of CV3-25 Fab in complex with the S2_1140-1165_ stem peptide. The overall structure of the complex is shown as a ribbon diagram (top panel) and with the molecular surface displayed over the Fab (bottom panel). S2 peptide (orange) assumes a bent conformation with the N-terminal α helical portion binding primarily to CDRs H1 and H2 (light green and cyan, respectively) and the random coil region interacting with CDR H3 (light pink). A non-canonical disulfide between C^99^ and C^100D^ of CDR H3 stabilizes the protruding hairpin in the CDR and likely strengthens its interactions with the C-terminal loop region of the S2 peptide. (**B**) Close-up views into the CV3-25 Fab-S2_1140-1165_ peptide interface. In the top panel the electrostatic potential (colored red, blue and white for negative, positive and neutral electrostatic potential respectively) is displayed over the molecular surface of the CV3-25 Fab (left) or S2-peptide (right) with 180° views of the complex. The bottom panel shows the network of H-bonds and salt bridges formed at the interface with orientations of complex as in the top panel. The putative glycosylation site at N^1158^ on the S2 peptide is marked with an asterisk. Salt bridges and hydrogen bonds with bond lengths < 3.5 Å as calculated by PISA^15^ (https://www.ebi.ac.uk/pdbe/pisa/) are denoted as yellow and blue dashed lines, respectively. A total of 11 H-bonds and 4 salt bridges are formed at the interface, with D^1153^ and K^1157^ of the S2 peptide contributing the majority of the hydrophilic contacts. The S2 bend and loop conformation (1158-1165) are stabilized by contacts to CDRs H1/H2/H3 with 4 H-bonds and 1 salt-bridge formed at the interface In addition, π-proline-π sandwich stacking interactions formed between the conserved residues F^1156^ and P^1162^ of S2 and Y^32^ of CDR H1 further stabilize the interface. CV3-25 light chain contacts are limited to only a single water mediated H-bond to the C-terminal D^1165^ of S2. (**C**) The network of interaction between residues at the CV3-25 Fab and the S2-peptide interface. Fab residues in the framework and CDRs of the Fab are colored as in (A) with interactions defined by a 5-Å distance criterion cutoff shown as lines. Salt bridges and H-bonds (bond length less than 3.5 Å) are shown as red dashed and blue solid lines, respectively. Hydrophobic interactions or bond distances between 3.5–5.0 Å are shown as grey dotted lines. (**D**) Diagram showing the buried surface area (BSA) of each individual S2 peptide residue in the CV3-25 Fab-S2 peptide complex. The BSA values of individual S2 residues were calculated using PISA^15^ (https://www.ebi.ac.uk/pdbe/pisa/) and are shown as the average of the values obtained for two complexes in the asymmetric unit of the crystal. (**E**) Sequence alignment of the S glycoprotein stem-peptide regions from representative beta-coronaviruses and two human alpha-coronaviruses. The helical regions as determined from PDB entry 6XR8 (residues 1140-1145) and the CV3-25-peptide structure (residues 1146-1156) are indicated above the sequence. S2 residues involved in the Fab-peptide interface are marked above the sequence with (*) and hydrogen-bonded or salt-bridged residues are marked with (+) for side chain, (-) for main chain and (±) for both side chain and main chain. The identical residues as compared to SARS-CoV-2 are highlighted in red with conservative changes marked in orange and non-conservative changes in black. (**F**) SPR sensorgrams of three SARS-CoV-2 S2 peptides binding to the immobilized CV3-25 IgG on a Protein A chip. The experimental data (colored) are fitted to a 1:1 Langmuir model (black) and the resulting kinetic constants are as shown. The minimal peptide recognized by CV3-25 with a low K_D_ value (∼1 μM) is 1153-DKYFKNHTSPD-1163. A 2 or 4 aa C-terminal extension leads to a 18 to 32 fold increase in the binding affinity (with a the 2-10 fold increase to the binding on-rate). (**G**) S2-peptide based structural comparison of CV3-25 Fab-SARS-CoV-2 S2_1140-1165_ peptide and P6-MERS-CoV S2_1230-1240_ peptide (PDB code: 7M55) in two orthogonal views. Only the variable regions of both Fabs are shown as surfaces for clarity.

To conceptualize the X-ray structure of the peptide bound to CV3-25 in the context of the S trimer, we superimposed two CV3-25 Fabs structures to the stem helix of the S trimer (PDB: 6XR8 (Cai et al., 2020) (Figure 7A). Direct superposition results in a clash of the Fabs and a mismatch of the Fab with the density map observed in the cryoET structure (Figure 7A). The random coil of the stem helix bound with CV3-25 points toward the center of the stem helix bundle producing a clash when two coils occupy the center region (Figure 7A). By performing flexible fitting we arrived at a structure for two CV3-25 Fabs bound to the S trimer, in which the helix and flexible turn are almost maintained at the original position for the first Fab (rotated by about 13°), and are shifted outward and rotated by about 20^°^ for the second Fab (Figure 7B, Video S2). The increasing need for dislocation and rotation likely explains that binding additional Fabs comes at an energy cost resulting in an asymmetric arrangement of one or two CV3-25 Fabs bound to S.

**Figure 7.**
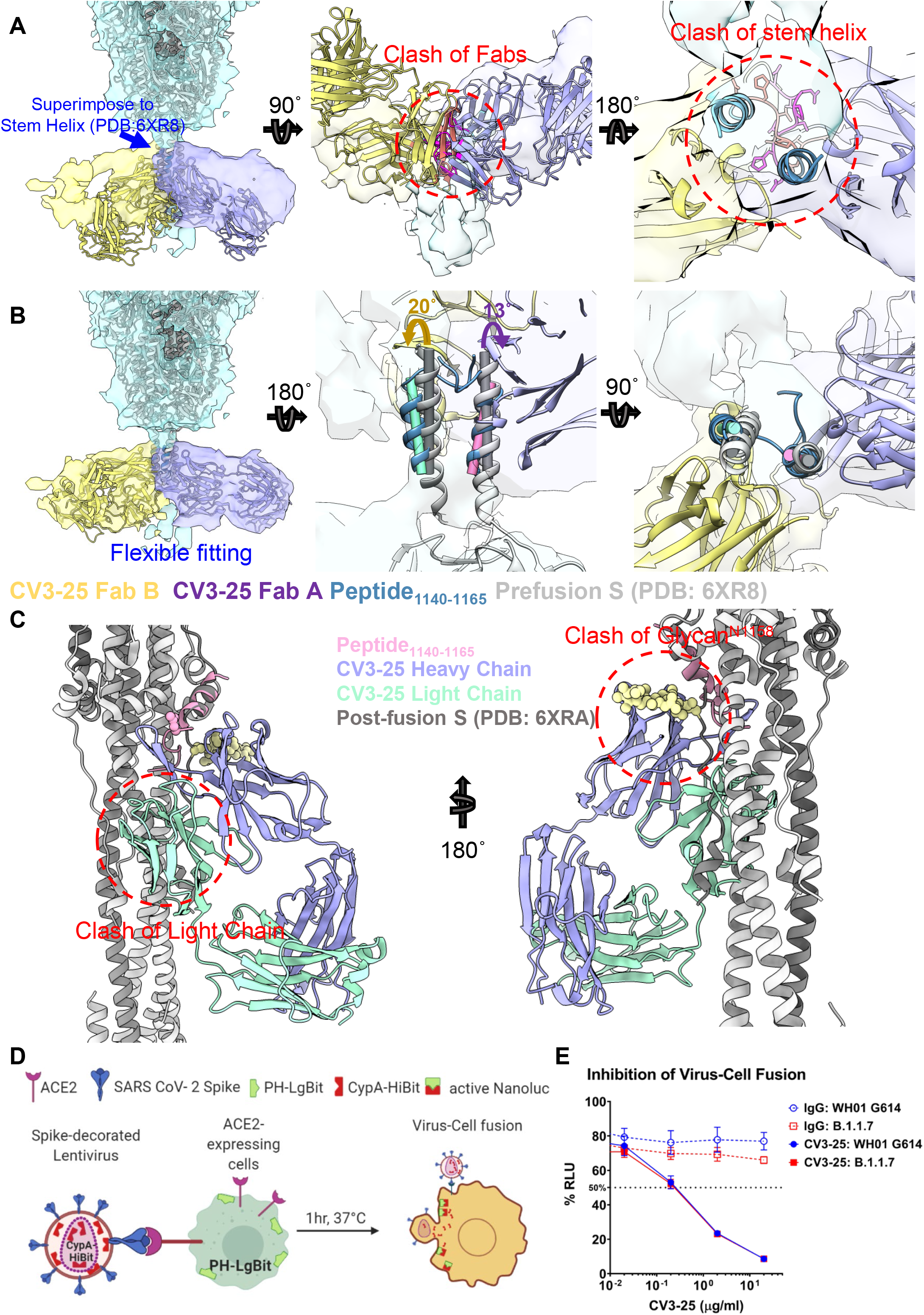
CV3-25 Inhibits S-Mediated Virus Fusion. (**A**) The crystal structures of the CV3-25 Fabs (purple and yellow) with the S2_1140-1165_ peptides (blue) were superimposed onto the stem helix of the prefusion S atomic model (gray, PDB 6XR8) and fitted into the CV3-25 CryoET structure. Left panel: side view indicates that CV3-25 does not dock into the cryoET density map. There are clashes between two Fabs (middle panel, bottom view) and the tails of binding stem helix (right panel, top view). (**B**) Flexible fitting with a combined model containing the CV3-25_S2_1140-1165_ crystal structure and prefusion S structure (gray, 6XR8) onto the cryoET structure. CV3-25 Fabs (purple and yellow) dock into the cryoET density map (left panel). The torsions of binding stem helix (blue, helix axes in cyan and pink, respectively) after fitting, comparing to original position of stem helix in 6XR8 (gray, helix axes in dark gray), were shown in middle and right panels. The third protomer was omitted for clarity. (**C**) Superimposition of peptide bound CV3-25 (purple, heavy chain; green, light chain) to the - fusion S (gray, PDB 6XRA). The peptide (pink) was aligned to the stem helix in the spike. The glycan on residue ASN1158 are shown in sphere representation (yellow). Possible clashes are indicated in red circles. (**D, E**) Investigation of Virus-Cell fusion activity in presence and absence of CV-35 mAb by the split nanoluc complementation assay. A scheme of the spilt nanoluc complementation assay experimental design was shown in (C).

Superimposition of the CV3-25/peptide structure to post-fusion S (PDB: 6XRA) (Cai et al., 2020) indicates that the light chain of CV3-25 would clash with HR1 of the adjacent protomer (Figure 7C). Furthermore, in post-fusion conformation, the stem helix unwinds a full turn, resulting in a clash between the glycan at ASN1158 and the heavy chain of CV3-25 (Figure 7C). These observations suggest that the structure of CV3-25 bound to S is incompatible with the post-fusion conformation of S. Indeed, we observed potent inhibition of membrane fusion by CV3-25 in a virus-to-cell fusion assay that uses nano-luciferase complementation (Figure 7D, E). As observed for CV3-1, CV3-25 also exhibits potent inhibitor function.

## Discussion

Here we describe the structures and mode of action of two potent anti-SARS-CoV-2 Spike antibodies. Both antibodies remained effective against emerging variants of concern and therefore were prime candidates to elucidate mode of action and identify epitopes with pan-coronavirus activity. CV3-1 stabilized the receptor-binding domain (RBD) in the “RBD-up” conformation and triggered potent shedding of S1. The ability of CV3-1 to neutralize variants of concern correlated with its ability to shed S1 and inactivate S. In contrast, CV3-25 bound to a highly conserved epitope in the stem helix in the S2 subunit and inhibited membrane fusion. We believe that both epitopes of these two antibodies are of interest for passive and active immunization strategies against emerging variants.

The cryoET structure of CV3-1 to S suggested binding to the 485-GFN-487 loop of RBD, an interpretation confirmed by mutagenesis. While mutations in these positions abrogate the binding of CV3-1 to S, they are rarely observed among circulating strains, suggesting that they are associated with a high fitness cost likely due to their importance in ACE2 interaction. Interestingly, CV3-1 exhibited potent agonist features indicating that it hits an allosteric site that is critical for the ability of ACE2 to induce conformational changes that lead to fusion. Consistent with this observation, CV3-1 induced potent shedding and the straightening of spikes indicative of allosteric signaling from the RBD all the way to the S2 stem region. This allosteric signaling likely weakens the S1-S2 interface leading to the observed shedding of S1. While CV3-1 is specific against SARS-CoV-2, it remained active against all tested variants of concern and variants of interest and protected K18-hACE2 transgenic mice from lethal challenges using the B.1.351 variant of concern. The potent agonist features within the ACE2 binding site may also open an opportunity for small molecule inhibitors that prematurely activate S not unlike CD4 mimetics in the case of HIV-1 envelope (Laumaea et al., 2020).

The structures of CV3-25 with a S peptide and intact S on the surface of virus particles revealed that it binds to the S2 stem in a region conserved among β-coronaviruses. Unlike other recently reported anti-S2 antibodies, CC40.8 and S2P6, which mainly recognize the stem helix and barely interact with the hinge region (Pinto et al., 2021; Zhou et al., 2021), CV3-25 also engages the hinge peptide known to be responsible for the tilting of spikes with respect to the membrane (Ke et al., 2020; Turoňová et al., 2020). This added ability of CV3-25 likely offers an advantage in capturing an easily accessible epitope in the hinge region and subsequently progressively twisting the helix to establish contact with the α-helical region of the stem. The relative conservation of this hinge is likely related to the observed allosteric communication from the RBD all the way down to S2. The post-fusion conformation of S forms a six-helix bundle structure when pulling two membranes together for fusion. This conformational change probably involves unwinding of the stem helix and loop to helix transition for the loose loop at the lower end. CV3-25 binding at both helix region and the random coil at the stem helix likely interrupts this S2 refolding, thus inhibiting membrane fusion.

One of the most exciting aspects of CV3-25 is its linear peptide epitope, which offers easy access to exploration of its potential as an immunogen. As the structure of the native S on the surface of virus particles revealed, access of CV3-25 is hindered by the need for rotation of the stem helix. However, such conformational readjustment is not needed for an immunogen. As such, eliciting antibodies targeting this S2 stem epitope using peptide or scaffold-presented peptide immunogens is predicted to be easier than when the entire spike trimer is the antigen. The potential of the CV3-25 epitope described herein should be explored as a candidate immunogen for vaccines that could be effective against all emerging variants and possibly exhibit pan-coronavirus efficacy.

## Acknowledgement

We thank Dr. Shenping Wu at Yale CryoEM facility for her technical assistance, and Dr. Zhuan Qin at University of Oxford for discussion on image processing. The authors thank the CRCHUM Animal Facility, BSL3 and Flow Cytometry Platforms for their technical assistance. We thank Dr. Stefan Pöhlmann and Dr. Markus Hoffmann (Georg-August University) for the plasmids coding for SARS-CoV-2 and Daniel Kaufmann for S2 15-mer peptides. The authors are grateful to MediMabs for providing their rabbit immunization protocol used to generate the anti-SARS-CoV-2 RBD polyclonal antibody. CV3-1 and CV3-25 antibodies were produced using the pTT vector kindly provided by the Canada Research Council. Crystallographic data were collected at the Stanford Synchrotron Radiation Light Source, SLAC National Accelerator Laboratory, which is supported by the U.S. Department of Energy, Office of Science, Office of Basic Energy Sciences, under contract number DE-AC02-76SF00515. The SSRL Structural Molecular Biology Program is supported by the DOE Office of Biological and Environmental Research and by the National Institutes of Health, National Institute of General Medical Sciences. This work was supported by a CIHR operating grant Pandemic and Health Emergencies Research/Project #465175 to M.P., W.M. and A.F., a NIH R01 AI163395-01 to W.M., by le Ministère de l’Économie et de l’Innovation (MEI) du Québec, Programme de soutien aux organismes de recherche et d’innovation to A.F., the Fondation du CHUM, a CIHR foundation grant #352417 to A.F., CIHR stream 1 and 2 for SARS-CoV-2 Variant Research to A.F. and M.C., an Exceptional Fund COVID-19 from the Canada Foundation for Innovation (CFI) #41027 to A.F., the Sentinelle COVID Quebec network led by the Laboratoire de Santé Publique du Quebec (LSPQ) in collaboration with Fonds de Recherche du Québec-Santé (FRQS) and Genome Canada – Génome Québec, and by the Ministère de la Santé et des Services Sociaux (MSSS) and MEI to A.F. A.F. and M.C. are recipients of Canada Research Chairs on Retroviral Entry no. RCHS0235 950-232424 and CRC in Molecular Virology and Antiviral Therapeutics, respectively. J.P. and S.P.A. are supported by CIHR fellowships, M.W.G. by the Gruber foundation, and R.G. by a MITACS Accélération postdoctoral fellowship. The funders had no role in study design, data collection and analysis, decision to publish, or preparation of the manuscript.

## Disclaimer

The views expressed in this presentation are those of the authors and do not reflect the official policy or position of the Uniformed Services University, U.S. Army, the Department of Defense, or the U.S. Government.

## Supplementary Figure Legends

**Figure S1.**
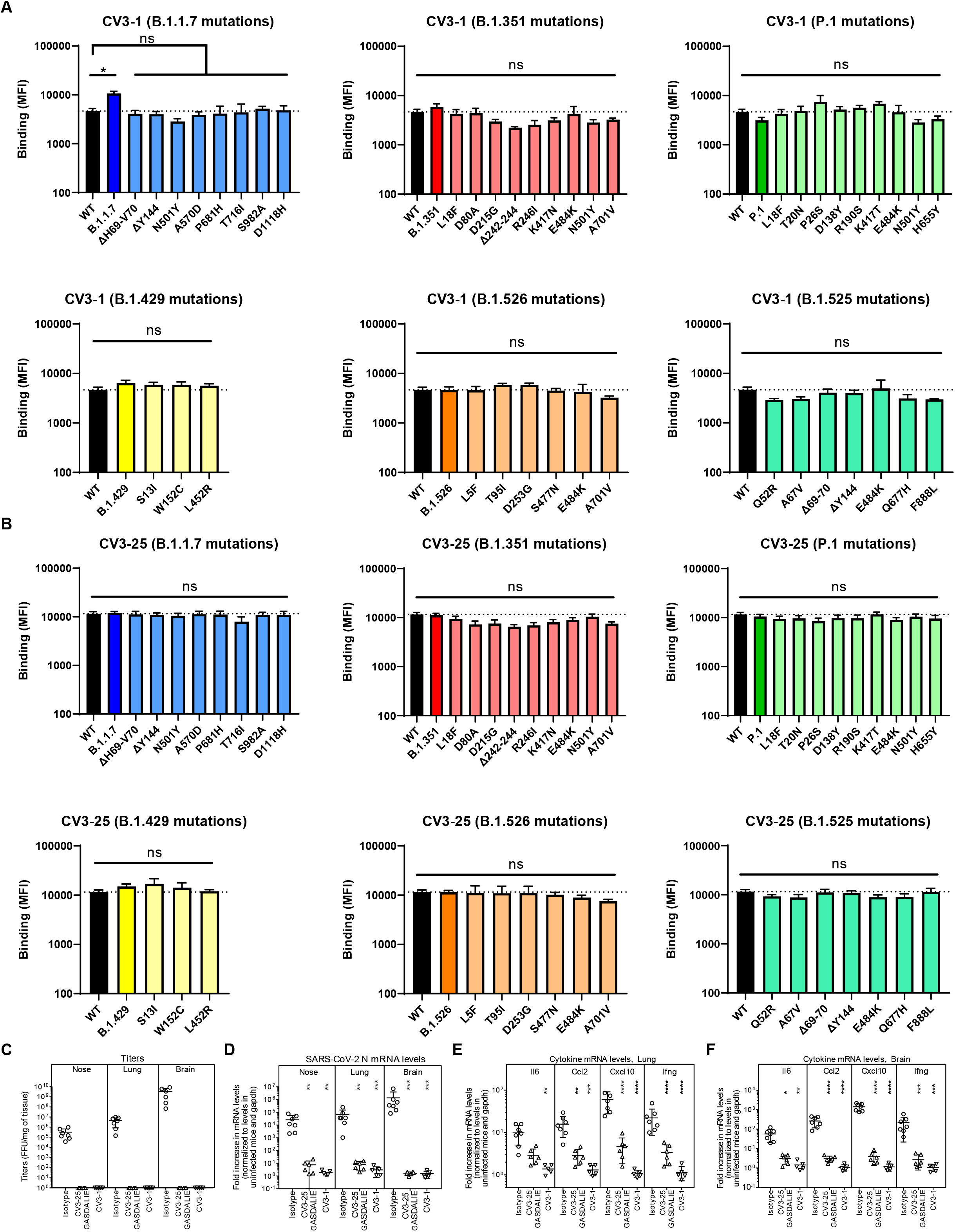
CV3-1 and CV3-25 Neutralize SARS-CoV-2 Variants In Vitro and Protect In Vivo. Related to Figure 1. (**A, B**) Cell-surface staining of 293T cells expressing full-length Spike from indicated variants (B.1.1.7, B.1.351, P.1, B.1.429, B.1.526, B.1.525) or their corresponding individual mutations by CV3-1 (A) and CV3-25 (B) mAbs. The graphs show median fluorescence intensities (MFIs). Dashed lines indicate the reference value obtained with Spike D614G. Error bars indicate means ± SEM. These results were obtained in at least two independent experiments. Statistical significance was tested using Kruskal-Wallis test with a Dunn’s post-test (*p < 0.05; ns, non significant). (**C**) Viral loads (FFUs/mg) from indicated tissue using Vero E6 cells as targets in mice prophylactically treated with CV3-1 and CV3-25 GASDALIE for the experiment shown in Figure 1C. Undetectable virus amounts were set to 1. (**D**) A plot showing mRNA levels SARS-CoV-2 nucleocapsid (N gene) from nose, lung and brain tissues of K18-hACE2 mice after sacrifice at times indicated in Figure 1E. (**E-F**) A plot showing mRNA levels of indicated cytokines from lung and brain tissues of K18-hACE2 mice after sacrifice at times indicated in Figure1E. The mRNA amounts in (D-F) were normalized to Gapdh mRNA and to levels seen in uninfected mice. Viral loads and inflammatory cytokine profile in indicated tissues were determined after necropsy for mice that succumb to infection at day 6 and for surviving mice at 10 dpi. Grouped data in (C-F) were analyzed by 2-way ANOVA followed by Tukey’s multiple comparison tests.

**Figure S2.**
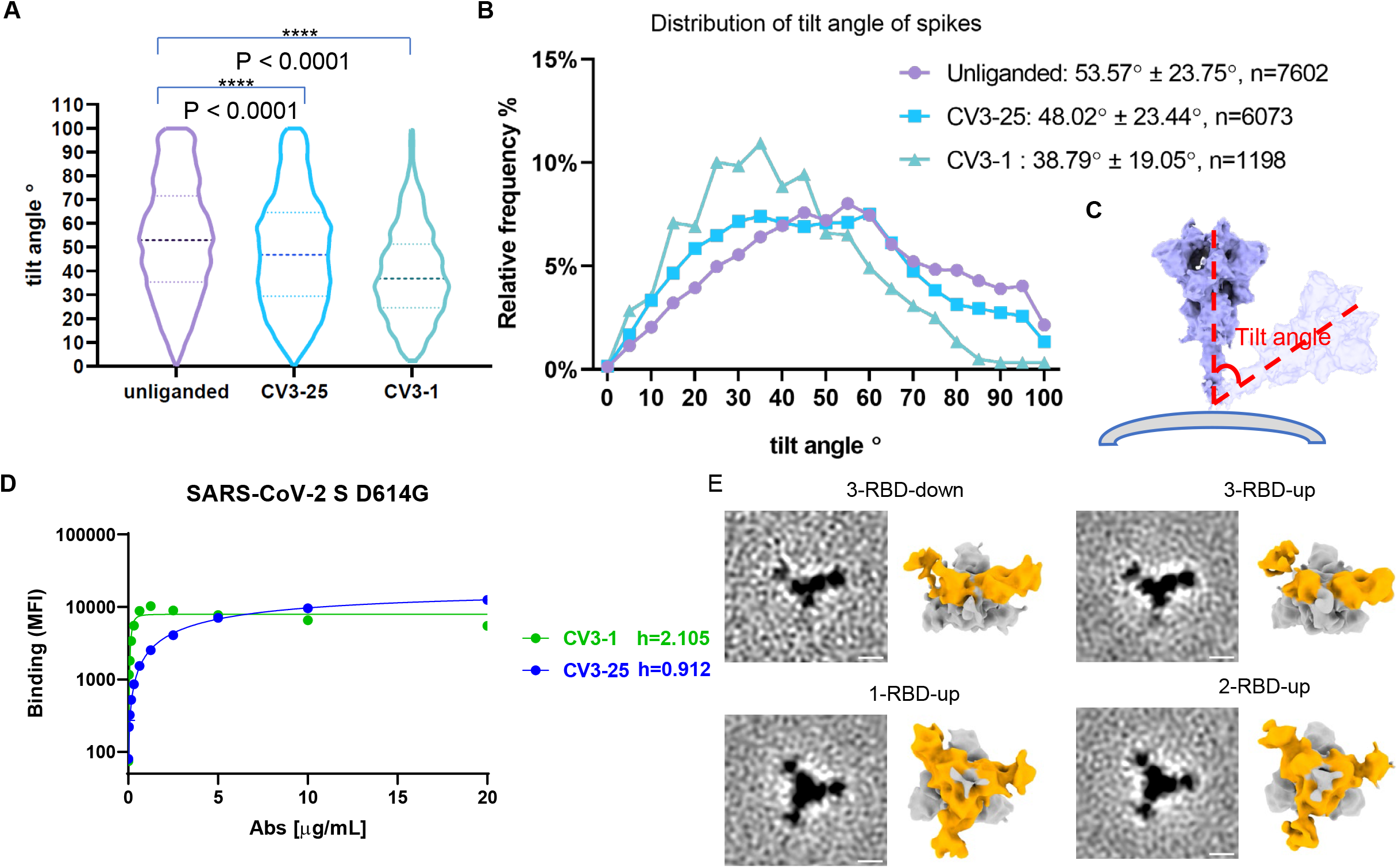
Conformational Dynamics of CV3-1 and CV3-25 Bound S_B.1.1.7_. Related to Figure 2. (**A-C**) Tilt angles of spikes on unliganded, CV3-1 Fab treated, and CV3-25 Fab treated pseudoviruses. Scheme graph of tilt angle is shown in (E). (**D**) The binding of CV3-1 or CV3-25 to SARS-CoV-2 S D614G expressed on 293T cells was measured flow cytometry. Cells were incubated with increasing amounts of mAbs and their binding was detected using a goat anti-human IgG AlexaFluor647. The Hill coefficients were determined using GraphPad software. These results were obtained in 3 independent experiments. (**E**) Subclass averages obtained after focused classification on the RBD of CV3-25 bound S. Bottom views (left) and segmentations (right) are shown for 3-RBD-down, 1-RBD-up, 2-RBD-up and 3-RBD-up classes. CV3-25 Fabs are shown in orange.

**Figure S3.**
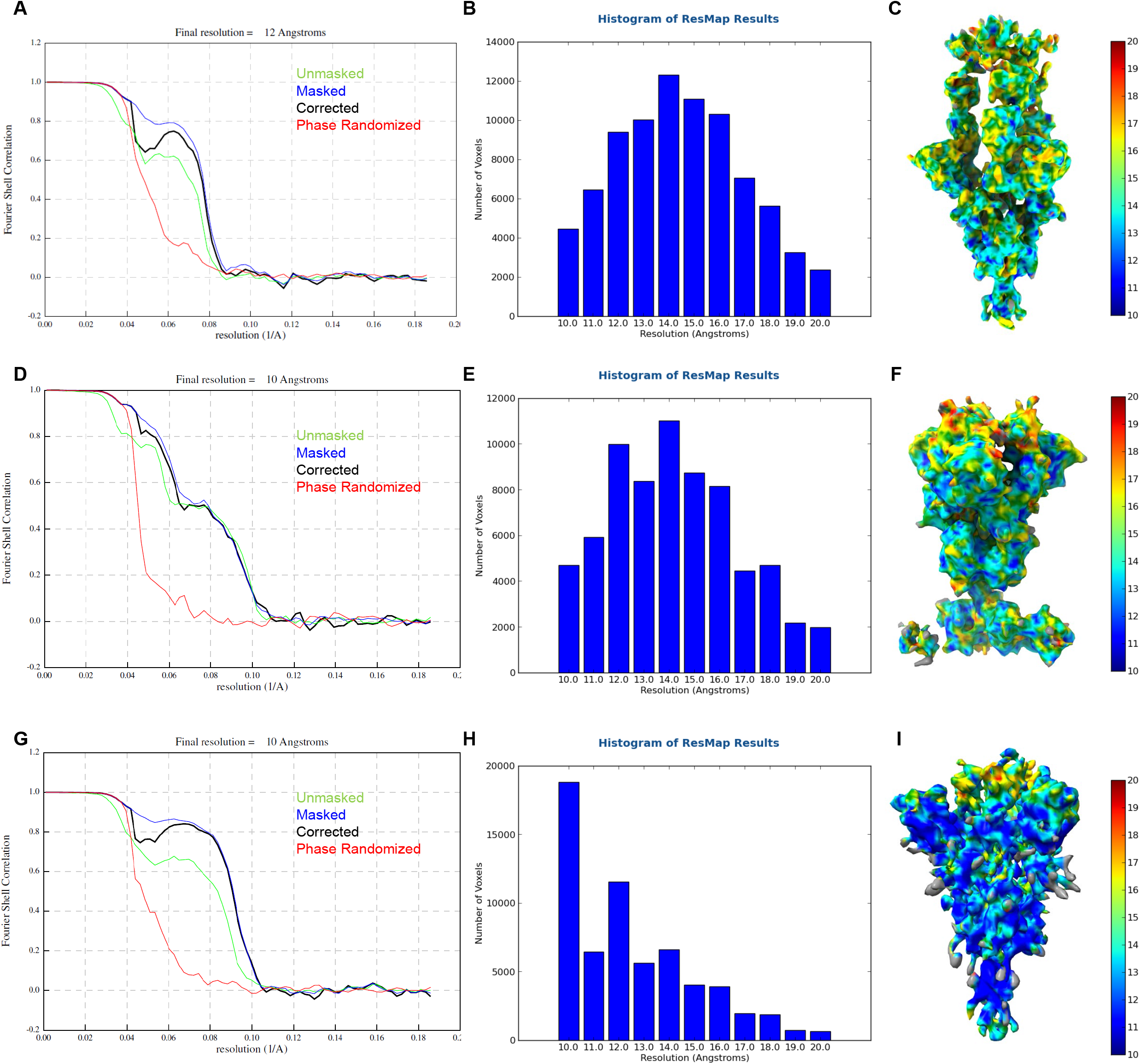
Resolution Assessment of Subtomogram Averaging Structure for CV3-1 Bound Spike. Related to Figures 3 and 5. (**A, D, G**) Resolution estimation based on Fourier shell correlation curves and 0.143 as a cutoff value. (**B, E, H**) Local resolution is estimated with Resmap. (**C, F, I**) Subtomogram averaged structures are colored according to the local resolution.

**Figure S4.**
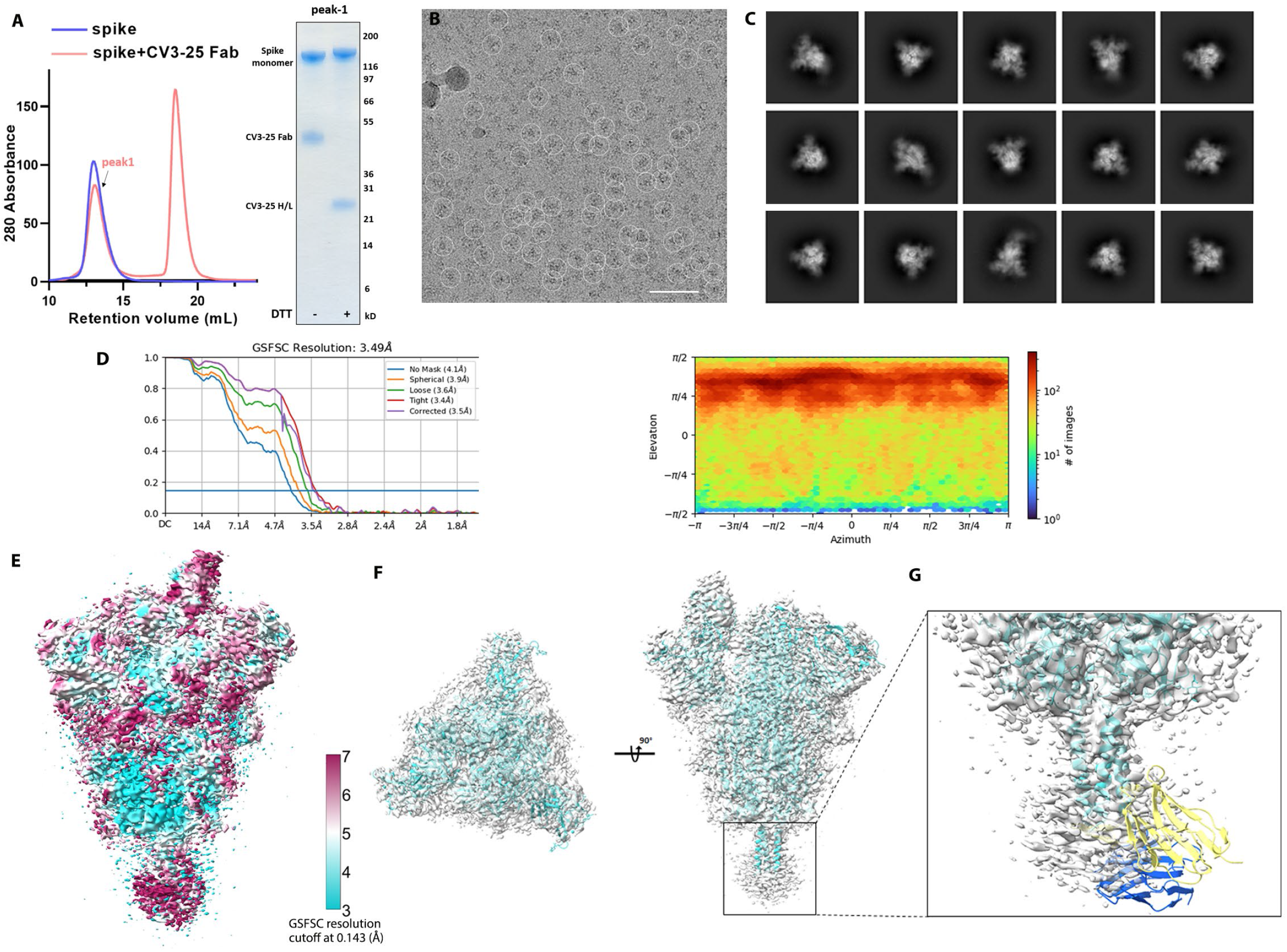
Cryo-EM Data for the Complex of CV3-25 Fab with SARS-CoV-2 HexaPro Spike. Related to Figure 5. (**A**) Cryo-EM sample preparation. Size-exclusion chromatogram of the purified, non-tagged SARS-CoV-2 HexaPro spike with CV3-25 Fab (molar-ratio: 1:20). SDS-PAGE analysis of peak1 of the spike Fab mixture shows that intact CV3-25 Fab is physically associated with the spike. (**B, C**) Representative electron micrograph after motion correction (B, scale bar 50 nm) and selected 2D averaged classes (C, in total 460k particles). (**D**) The Fourier shell correlation curves indicate an overall resolution of 3.49 Å using non-uniform refinement with C1 symmetry (left panel). The view direction distribution plot of all particles used in the final refinement shown as a heatmap (right panel). (**E**) The final overall map is shown and colored according to the local resolution as calculated in cryoSPARC using a FSC cutoff of 0.143. (**F**) Side and top views of the cryo-EM density map (semi-transparent grey surface) fitted with a prefusion spike model with a one-RBD-up conformation shown in cyan. An initial model template was generated using the NTD (residues 12-305) from PDB entry 7LY31, the RBD (residues 306-541) and S1-S2 core (residues 542-1139) from 6XKL, and the S2 stem helix (1140-1162) from 6XR8 with the “fit-in-map” function in chimeraX. (**G**) A S2-stem-peptide based superimposition of the variable region from the CV3-25-peptide crystal structure (yellow and blue) with the cryo-EM model mimics the one-Fab-bound state. The discrete, feeble and nearly-isotropic density around the S2-helix indicates that there is a high degree of local dynamic motion and a diverse collection of Fab-stem-peptide conformations/orientations relative to the rigid S2 core that may transiently coexist.

**Figure S5.**
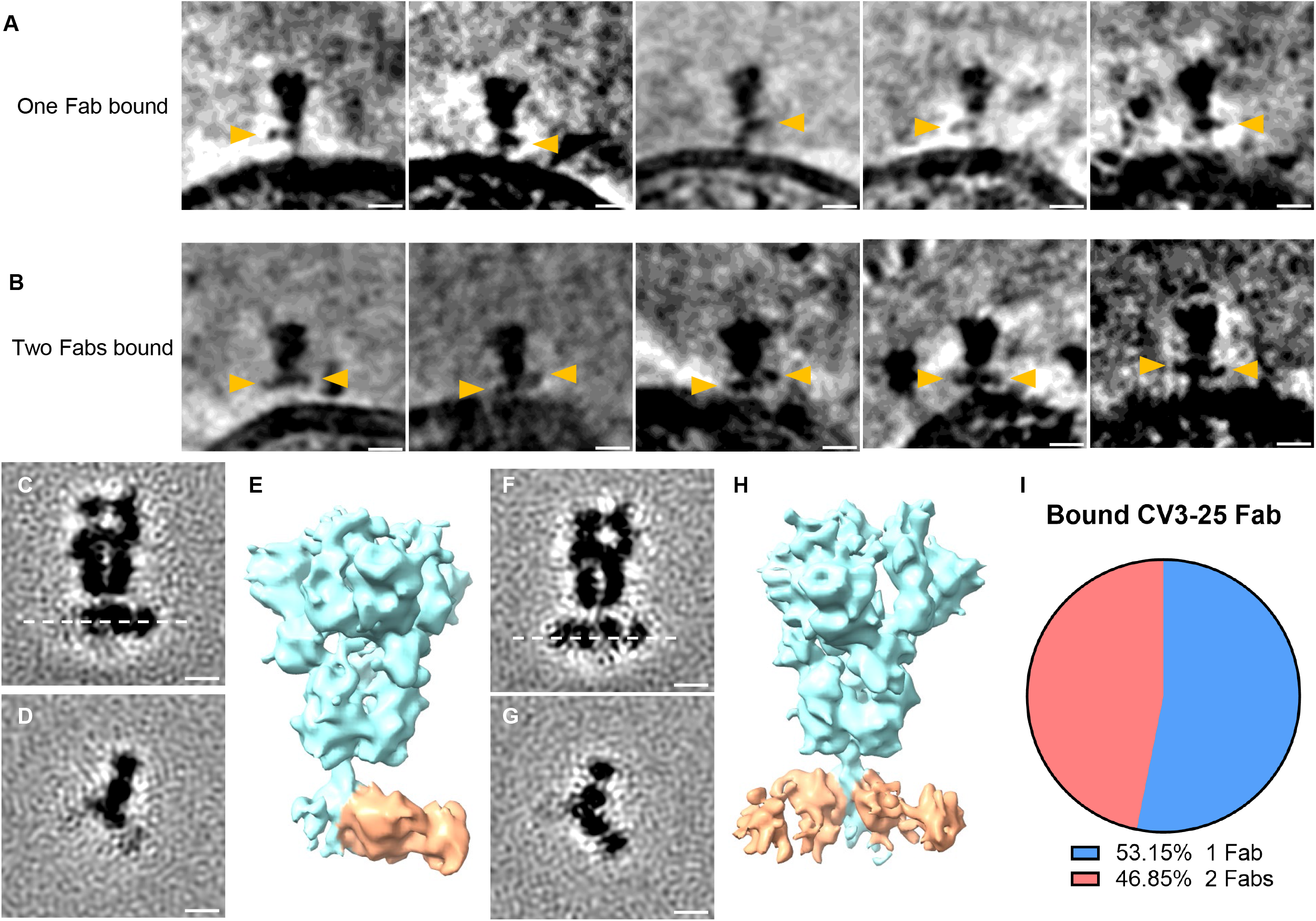
CV3-25 Binds on a Conserved Epitope on S2. Related to Figure 5. (**A-B**) Gallery of spikes bound to one CV3-25 Fab (A) and two CV3-25 Fabs (B) on lentiviral particles. CV3-25 Fabs are indicated by yellow arrowheads. (**C-H**) Side view (C, F) and top view (D, G) of averaged structure of S bound with one CV3-25 Fab (C-E) and two CV3-25 Fabs (F-H). Segmentations of the structures are shown in (E, H). CV3-25 Fabs are shown in orange and S is shown in cyan. (**I**) Proportion of S bound with one and two CV3-25 Fabs.

**Figure S6.**
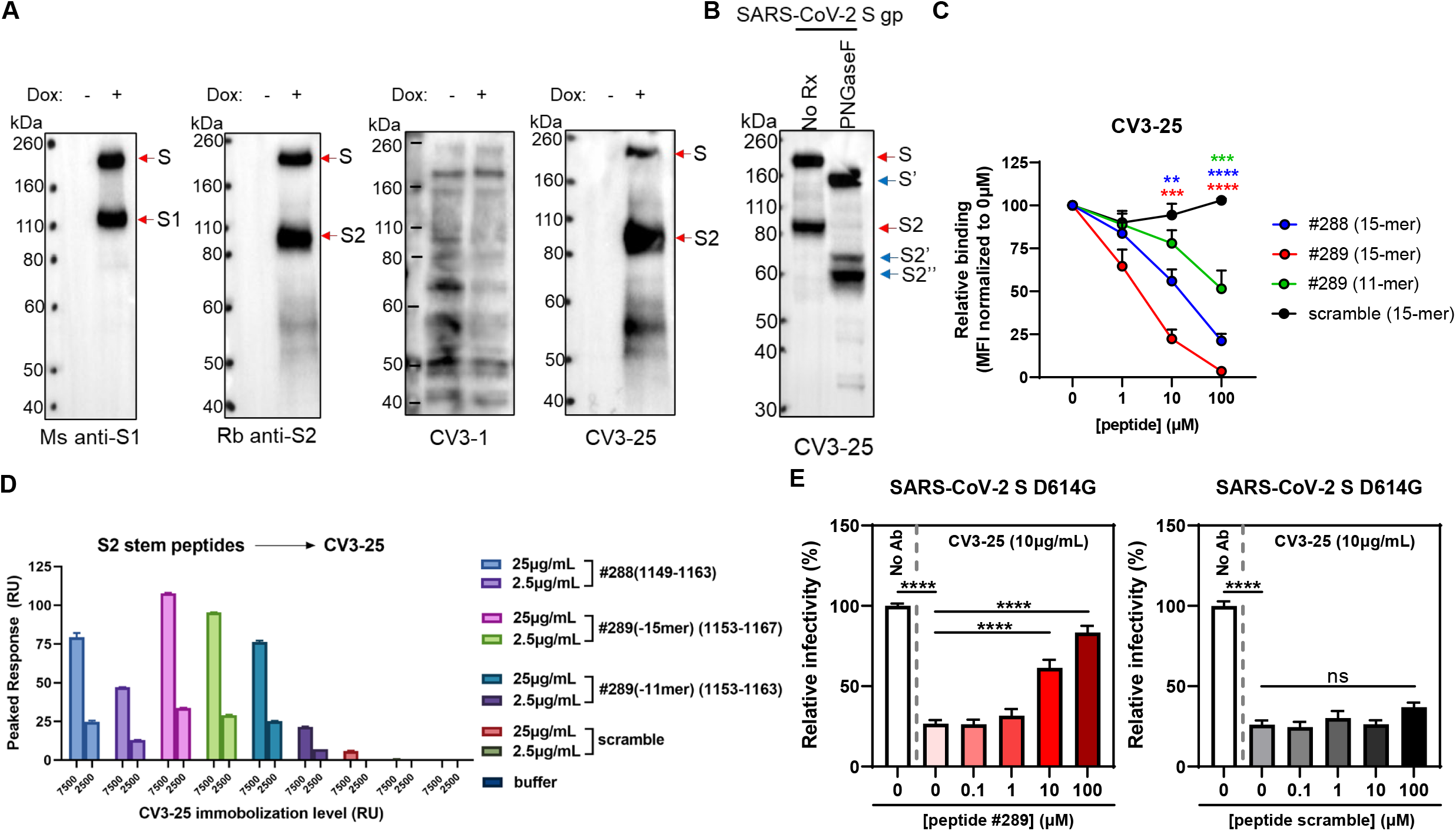
CV3-25 Binds on a Conserved Epitope on S2. Related to Figures 5 and 6. (**A**) 293T-S cells were induced with doxycycline (Dox) to express wild-type SARS-CoV-2 S glycoprotein, or mock treated as a control. Two days after induction, cells were lysed, and cell lysates were subjected to Western blotting with CV3-1 or CV3-25, or with a mouse (Ms) anti-S1 antibody or rabbit (Rb) anti-S2 antibody as controls. (**B**) 293T-S cells induced to express the SARS-CoV-2 S glycoprotein (gp) were lysed with lysis buffer. Cell lysates were then treated with PNGase F or, as a control, mock treated (No Rx). The cell lysates were then Western blotted with the CV3-25 antibody. S’, S2’ and S2’’ are deglycosylated forms of S or S2. (**C**) Cell-surface staining of 293T cells expressing the wild-type SARS-CoV-2 Spike with CV3-25 mAb in presence of increasing concentrations of S2 peptides #288 (15-mer), #289 (15-mer), #289 (11-mer) or a scrambled peptide (15-mer) as a control. The graphs show the median fluorescence intensities (MFIs) normalized to the condition without any peptide (0 µM). Error bars indicate means ± SEM. These results were obtained in 3 independent experiments. (**D**) SPR binding of SARS-CoV-2 S2 peptides to immbolized CV3-25. CV3-25 IgG was immobilized on a protein A chip either to 2500 or 7500 RU prior to peptide injection. The S2 peptides of different truncations or the equivalent scrambled peptides were injected at the indicated concentrations. The peak response was taken to be the background corrected response at the steady state where the binding reached equilibrium. The data shown is from three independent experiments. (**E**) Pseudoviruses encoding for the luciferase reporter gene and bearing SARS-CoV-2 Spike D614G were used to infect 293T-hACE2 target cells. Pseudovirions were incubated with CV3-25 mAb (10 µg/mL) in presence of increasing concentrations of S2 peptide #289, or peptide scramble as a control, for 1h at 37°C prior infection of 293T-hACE2 cells for 48h at 37°C. Error bars indicate means ± SEM. These results were obtained from at least 3 independent experiments. Statistical significance was tested using (C) one-way ANOVA with a Holm-Sidak post-test or (E) an unpaired T test (**p < 0.01; ***p < 0.001; ****p < 0.0001; ns, non significant).

**Figure S7.**
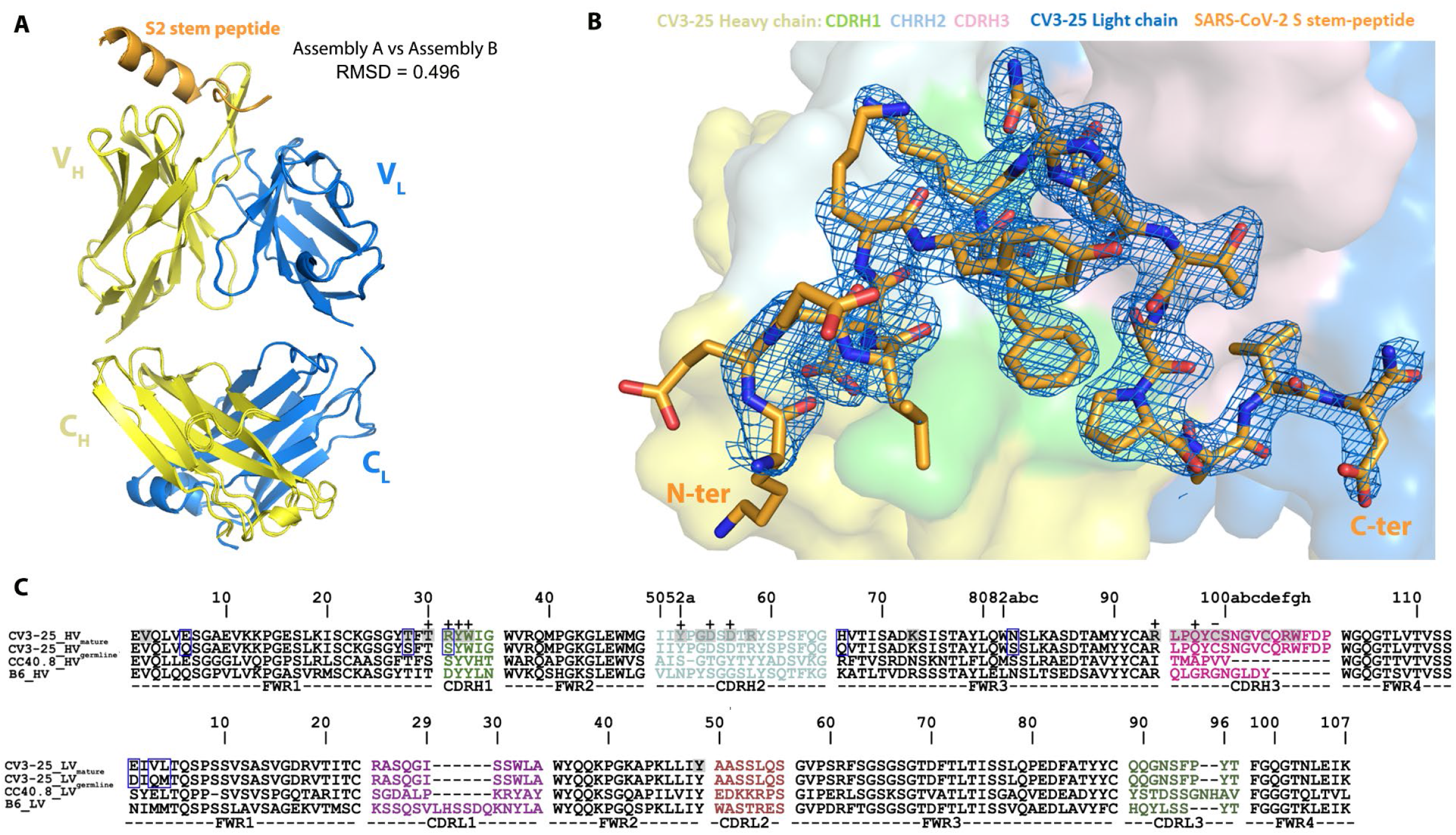
Crystal Structure of CV3-25-S2 Peptide Complex. Related to Figure 6. (**A**) Ribbon diagram of the superposition of the two copies of CV3-25 Fab-S2_1140-1165_ peptide complex from the asymmetric unit of the crystal. The overall structures of these two copies are very similar with a root mean square deviation (RMSD) of equivalent Cα atoms of 0.496 Ǻ for the complex, 0.499 Ǻ for the Fab and 0.305 Ǻ for the S2 peptide. (**B**) 2Fo-Fc electron density map of S2 peptide contoured at 1.5 σ The N-terminal non-contact S2 residues D^1146^ to K^1148^ are omitted for clarity. The surface of CV3-25 Fab is represented with CDR H1, CDR H2, and CDR H3 colored in green, cyan and pink respectively. (**C**) V_H_ and V_L_ sequence alignments of affinity matured CV3-25 and its germline IGHV5-51 along with two other S2-binding Abs, CC40.8 (anti SARS-CoV-2)^6, 17^ and B6 (anti MERS-CoV)^5^. CDR sequences are colored as indicated in Figure 6. The buried surface residues (BSA > 0 Å) as calculated by PISA^15^ are shaded in grey. Contact residues involved in salt-bridges or H-bonds to S2 peptide are marked above the sequence with (+) for the side chain and (-) for the main chain. Somatically mutated residues as compared to the germline sequence are highlighted with blue boxes. The peptide binding paratope of CV3-25 is predominantly formed by CDRs of the heavy chain which contributes ∼95% of the overall S2 buried surface area (BSA) from the Fab (BSA of 637.6 Å^2^ and 27.6 Å^2^ for the heavy and light chain, respectively). The paratope-epitope interface is comprised mostly of hydrophilic interactions which includes an extensive network of charge complementarity (Fig. 6B and C). In the α helical segment of the peptide there are 5 hydrogen-bonds (H-bonds) and 3 salt-bridges formed between the conserved S2 residue D^1153^ and K^1157^ (either side chain or backbone) with the CDR H1/H2 side chain atoms of residues T^30^, R^31^/D^54^, D^56^ or the main chain atoms of CDR H1/H2/H3 (residues W^33^/Y^52^/Q^97^) with the bonding distances < 3.2 Å. As a result, the highly conserved D^1153^ and K^1157^ have the highest BSA among the 20 structurally resolved S2 residues, underlying the structural basis of CV3-25’s broad neutralizing capacity against beta-coronaviruses.

Video S1. Flexible Fitting of CV3-1 Bound SARS-CoV-2 Spike cyroET Structure with 3-RBD-down atomic model (PDB 7lws). Both side view and top view were recorded.

Video S2. Flexible Fitting of 2 CV3-25 Bound SARS-CoV-2 Spike cyroET Structure with prefusion S (6xr8) superimposed with two CV3-25 models at the stem helix.

**Table S1.**
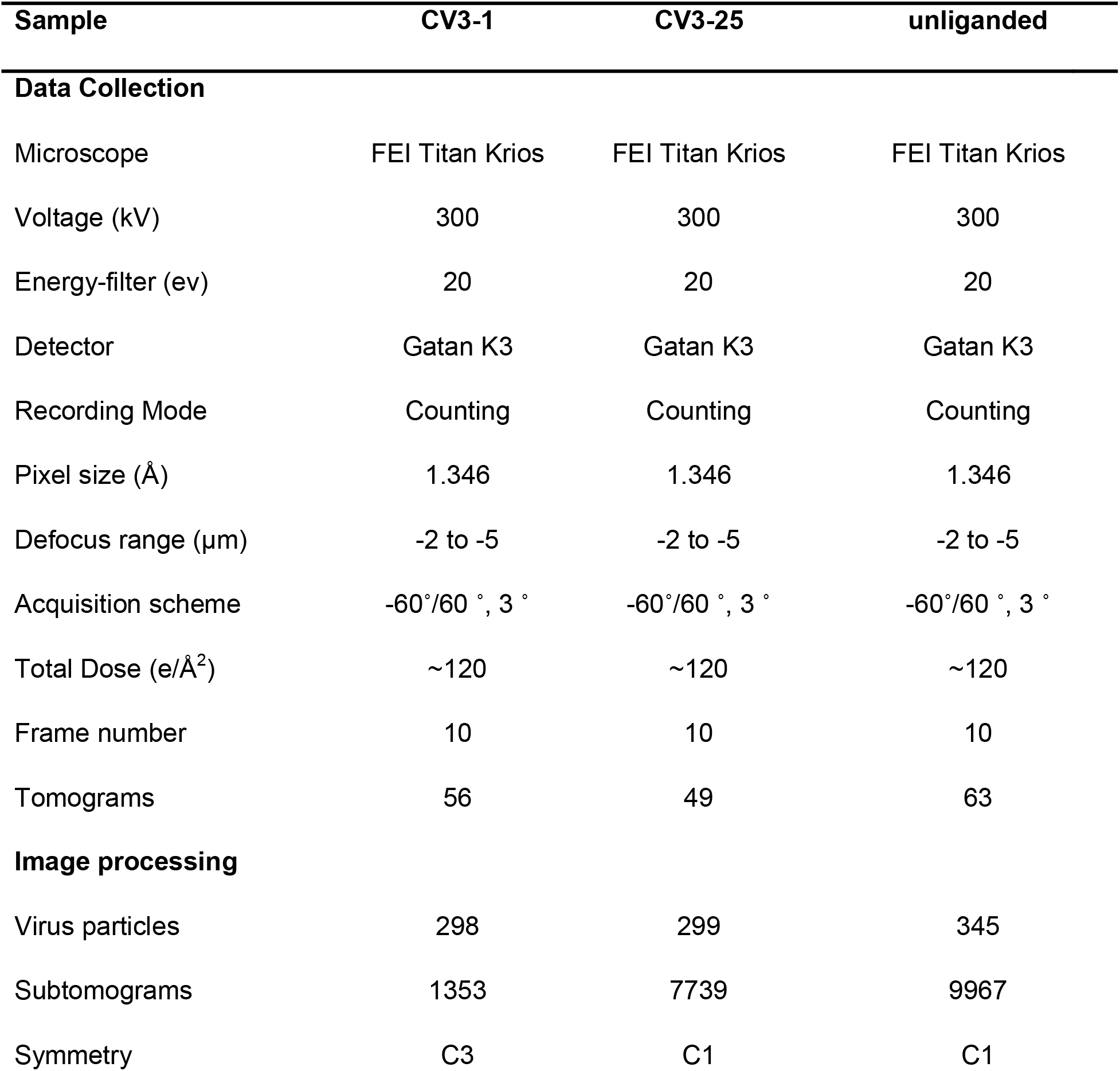
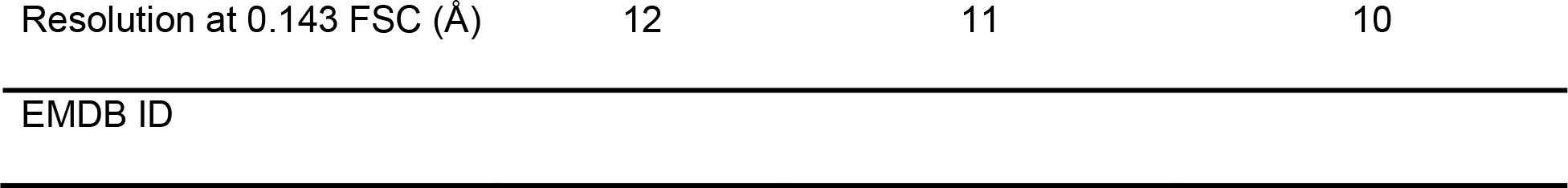
CryoET Data Acquisition and Image Processing.

**Table S2.** Crystallographic Data Dollection and Refinement Statistics.

| CV3-25 Fab_ S2<sub>1140-1165</sub> peptide complex |  |
| --- | --- |
| **Data collection** |  |
| Wavelength, Å | 0.979 |
| Resolution range, Å | 38.6 - 2.15 (2.23 - 2.15) |
| Space group | P2<sub>1</sub> |
| Unit cell parameter |  |
| a, b, c, Å | 82.8, 85.2, 87.1 |
| α, β, γ, ° | 90, 114.77, 90 |
| Redundancy | 25.7 (2.0) |
| Completeness, % | 96.5 (81.9) |
| Mean I/sigma(I) | 6.18 (2.3) |
| R<sub>merge</sub><sup>a</sup> | 0.185 (0.489) |
| R<sub>pim</sub><sup>b</sup> | 0.119 (0.329) |
| CC<sub>1/2</sub><sup>c</sup> | 0.942 (0.714) |
| Wilson B<sub>factor</sub>, (1/Å<sup>2</sup>)<sup>d</sup> | 38.7 |
| **Refinement** |  |
| R<sub>work</sub><sup>e</sup> | 0.196 (0.256) |
| R<sub>free</sub><sup>f</sup> | 0.239 (0.290) |
| Resolution, Å | 38.6 - 2.15 |
| # of non-hydrogen atoms |  |
| proteins | 7,032 |
| water | 530 |
| Overall B<sub>factor</sub>, (Å<sup>2</sup>) |  |
| proteins | 45 |
| ligands | 58 |
| water | 47 |
| RMS (bond lengths), Å | 0.009 |
| RMS (bond angles), ° | 1.17 |
| Ramachandran<sup>g</sup> |  |
| Favored, % | 97.8 |
| Allowed, % | 2.2 |
| Outliers, % | 0.0 |
| PDB ID | 7NAB |

Statistics for the highest-resolution shell are shown in parentheses.
<sup>a</sup> $R_{\text{merge}} = \sum |I - \langle I \rangle| / \sum I$ , where $I$ is the observed intensity and $\langle I \rangle$ is the average intensity obtained from multiple observations of symmetry-related reflections after rejections
<sup>b</sup> $R_{\text{pim}} =$ as defined in (Weiss, 2001)
<sup>c</sup> $CC_{1/2} =$ as defined by Karplus and Diederichs (Karplus and Diederichs, 2012)
<sup>d</sup>Wilson $B_{\text{factor}}$ as calculated in (Popov and Bourenkov, 2003)
<sup>e</sup> $R = \sum \|F_o\| - \|F_c\| / \sum \|F_o\|$ , where $F_o$ and $F_c$ are the observed and calculated structure factors, respectively
<sup>f</sup> $R_{\text{free}} =$ as defined by Brünger (Brunger, 1997)
<sup>g</sup>Calculated with MolProbity

**Table S3.**
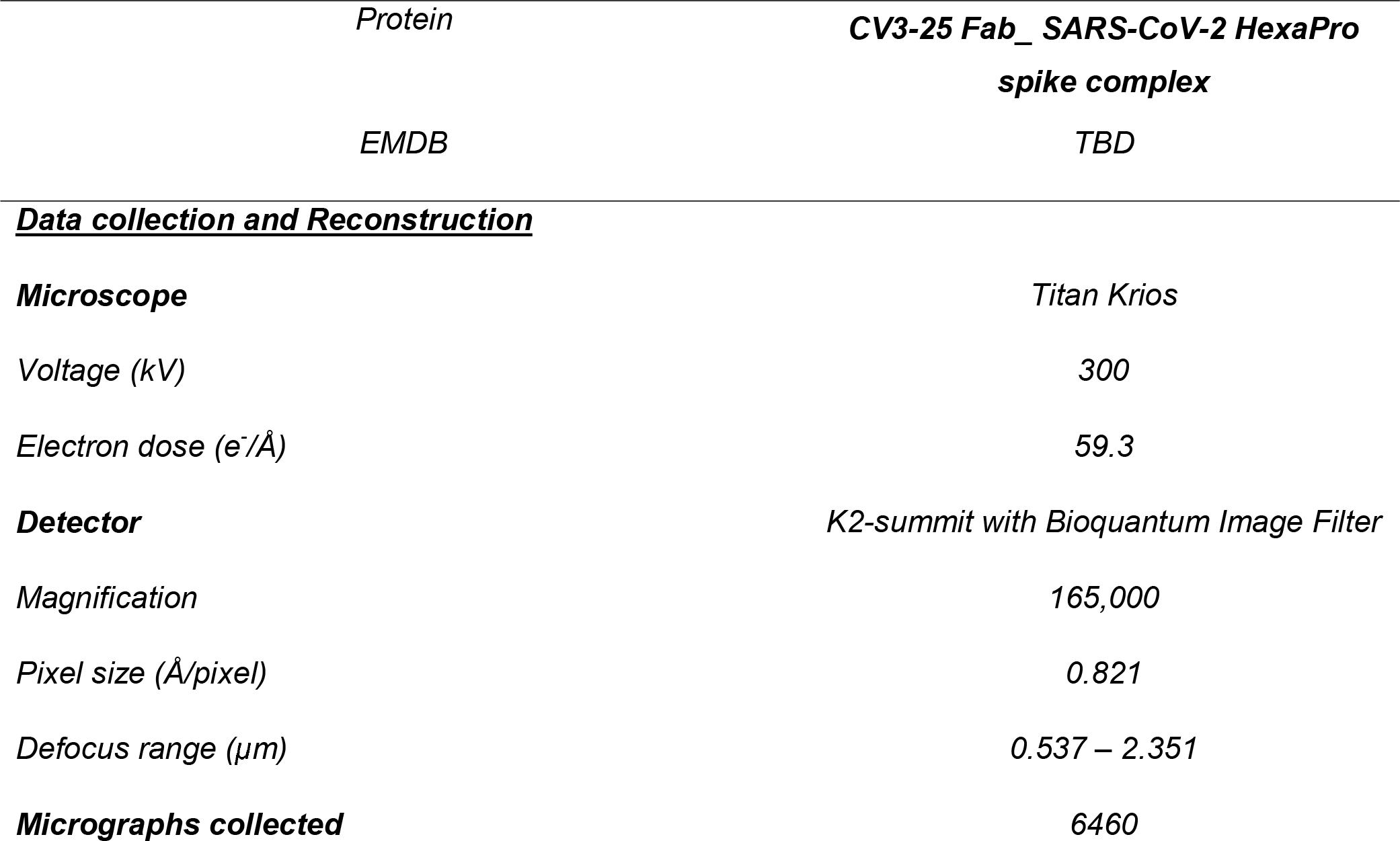
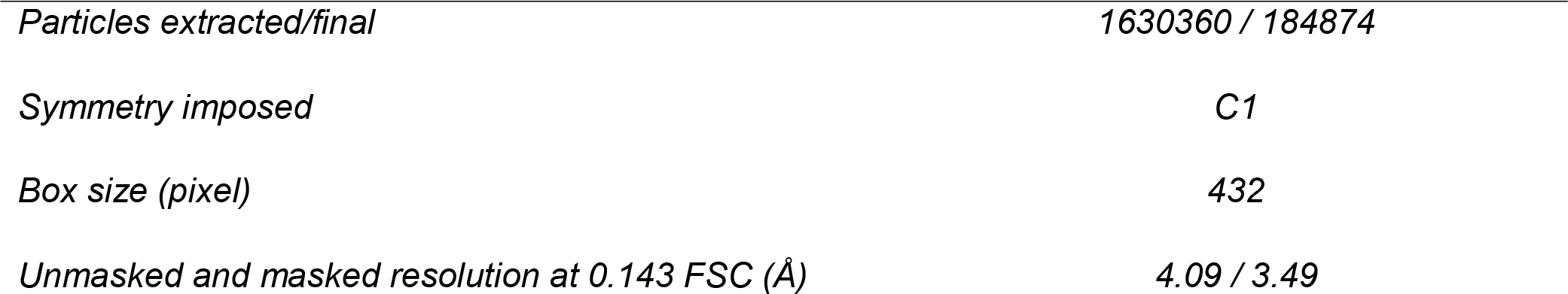
Cryo-EM Data Collection and Refinement Statistics.

| Protein | CV3-25 Fab_ SARS-CoV-2 HexaPro spike complex |
| --- | --- |
| EMDB | TBD |
| **<u>Data collection and Reconstruction</u>** |  |
| **Microscope** | Titan Krios |
| Voltage (kV) | 300 |
| Electron dose (e<sup>-</sup>/Å<sup>2</sup>) | 59.3 |
| **Detector** | K2-summit with Bioquantum Image Filter |
| Magnification | 165,000 |
| Pixel size (Å/pixel) | 0.821 |
| Defocus range (μm) | 0.537 – 2.351 |
| **Micrographs collected** | 6460 |
| *Particles extracted/final* | *1630360 / 184874* |
| *Symmetry imposed* | *C1* |
| *Box size (pixel)* | *432* |
| *Unmasked and masked resolution at 0.143 FSC (Å)* | *4.09 / 3.49* |

## METHOD DETAILS

### Cell Lines

*293T human embryonic kidney cells (ATCC) and 293T-ACE2 cells were maintained at 37°C under 5% CO_2_ in Dulbecco’s Modified Eagle Medium (DMEM) (Wisent), supplemented with 5% fetal bovine serum (FBS) (VWR) and 100 U/mL penicillin/streptomycin (Wisent). 293T-ACE2 cells stably expressing human ACE2 are derived from 293T cells and were maintained in medium supplemented with 2 µg/mL of puromycin (Millipore Sigma) (Prévost et al., 2020)*

### Plasmids and Site-Directed Mutagenesis

*The plasmids expressing the wildtype SARS-CoV-2 Spike was previously reported (Hoffmann et al., 2020). The plasmid encoding for SARS-CoV-2 S RBD (residues 319-541) fused with a hexahistidine tag was previously described (Beaudoin-Bussières et al., 2020). The individual mutations in the full-length SARS-CoV-2 Spike expressor, the furin cleavage site mutations (R682S/R683S) and the Spike from the B.1.429 lineage (S13I, W152C, L452R, D614G) were generated using the QuikChange II XL site-directed mutagenesis kit (Agilent Technologies). The amino acid deletions in the full-length SARS-CoV-2 Spike expressor were generated using the Q5 site-directed mutagenesis kit (NEB). The presence of the desired mutations was determined by automated DNA sequencing. The plasmids encoding the Spike from the B.1.1.7 lineage (*Δ*69-70,* Δ*144, N501Y, A570D, D614G, P681H, T716I, S982A and D1118H), the B.1.351 lineage (L18F, D80A, D215G,* Δ*242-244, R246I, K417N, E484K, N501Y, D614G, A701V), the P.1 lineage (L18F, T20N, P26S, D138Y, R190S, K417T, E484K, N501Y, D614G, H655Y, T1027I) and the B.1.526 lineage (L5F, T95I, D253G, E484K, D614G, A701V) were codon-optimized and synthesized by Genscript. The plasmids encoding the Spike from the B.1.617.1 (E154K, L452R, E484Q, D614G, P681R) and the B.1.617.2 (T19R, Δ156-158, L452R, T478K, D614G, P681R, D950N) lineages were generated by overlapping PCR using a codon-optimized wild-type SARS-CoV-2 Spike gene that was synthesized (Biobasic, Markham, ON, Canada) and cloned in pCAGGS as a template. All constructs were validated by Sanger sequencing. The plasmid encoding for the ACE2-Fc chimeric protein, a protein composed of an ACE2 ectodomain (1–615) linked to an Fc segment of human IgG1 was previously reported (Anand et al., 2020)*.

### Antibodies

*The human antibodies (CV3-1 and CV3-25) used in the work were isolated from the blood of convalescent donor S006 (male) recovered 41 days after symptoms onset using fluorescent recombinant stabilized Spike ectodomains (S2P) as probes to identify antigen-specific B cells as previously described (Jennewein et al., 2021). Site-directed mutagenesis was performed on plasmids expressing CV3-25 antibody heavy chain in order to introduce the GASDALIE mutations (G236A/S239D/A330L/I332E) using the QuickChange II XL site-directed mutagenesis protocol (Stratagene) (Ullah et al., 2021). Two New Zealand White rabbits were immunized with purified recombinant SARS-CoV-2 RBD proteins using MediMabs’ 77-day Canadian Council on Animal Care (CCAC)-accredited protocol. Animals were hosted and handled at the CRCHUM Animal Facility and the experimental protocol received approval from the Institutional Animal Protection Committee prior the beginning of the manipulation (protocol #IP18039AFl). The first immunization was done using complete Freund’s adjuvant (Millipore Sigma) followed by 4 immunizations with incomplete Freund’s adjuvant (Millipore Sigma). Rabbits were used solely for this project and were sacrificed by total exsanguination. Blood was processed and serum was further used in immunoprecipitation experiments at 1:1000 dilution*.

### Cryo-electron Tomography Sample Preparation

*Lentiviral particles were collected and clarified by low-speed spinning (1500g for 5 min) twice, then pelleted by ultracentrifugation (130,000g for 2 hour) once and resuspended in PBS buffer. 6 nm gold tracer was added to the concentrated S-decorated HIV-1 lentivirus at 1:3 ratio, and 5 µl of the mixture was placed onto freshly glow discharged holey carbon grids (R 2/1, Quantifoil) for 1 min. Grids were blotted with filter paper, and plunge frozen into liquid ethane by a homemade gravity-driven plunger apparatus. Frozen grids were stored in liquid nitrogen until imaging*.

### Cryo-electron Tomography Data Collection

*Cryo-grids were imaged on a cryo-transmission electron microscope (Titan Krios, Thermo Fisher Scientific) operated at 300 kV, using a Gatan K3 direct electron detector in counting mode with a 20 eV energy slit. Tomographic tilt series between −60° and +60° were collected by using SerialEM (Mastronarde, 2005)(Mastronarde, 2005) in a dose-symmetric scheme (Hagen et al., 2017; Mastronarde and Held, 2017)(Hagen et al., 2017) with increments of 3°. The nominal magnification was 64,000 X, giving a pixel size of 1.346 Å on the specimen. The raw images were collected from single-axis tilt series with accumulative dose of ∼120e per Å^2^. The defocus range was −2 to −6 µm and 9 frames were saved for each tilt angle. Detailed data acquisition parameters are summarized in* Table S1.

*Frames were motion-corrected using Motioncorr2 (Zheng et al., 2017)(Zheng et al.,2017) to generate drift-corrected stack files, which were aligned using gold fiducial makers by IMOD/etomo (Mastronarde and Held, 2017)(Mastronarde and Held, 2017). The contrast transfer function (CTF) was measured by the ctfplotter package within IMOD. Tilt stacks were CTF-corrected by ctfphaseflip within IMOD. Tomograms were reconstructed by weighted back projection and tomographic slices were visualized with IMOD*.

### Cryo-electron Tomography Data Analysis

*For the CV3-1 sample, all spikes were manually picked. Eular angles were determined based on the vector between two points, one on the head of the spike and the other on the membrane where the spike locates. For CV3-25 and unliganded samples, a low-pass filtered (30Å) structure from previous aligned S structure was used as the template for template matching search in 8 x binned tomograms. Subtomograms were extracted for initial alignment. After this alignment, particles with cross-correlation coefficients (CCC) below 0.25 were removed. Visual inspection of the tomograms in IMOD confirmed that the rest of the subtomograms corresponded to S trimers on the viral surface. Particles that had tilted by more than 90° relative to their perpendicular positions to the viral surface were excluded. Subsequent processing was performed by using I3 (Winkler, 2007) with 2 x and 4 x binned tomograms*.

*All the density maps were segmented in the UCSF Chimera (Pettersen et al., 2004), and ChimeraX (Goddard et al., 2018; Pettersen et al., 2021) was used for surface rendering and visualization of cryo-ET maps and models. “Fit in map” tool in Chimera and ChimeraX was used for rigid fitting. iMODFIT was used for flexible fitting (Lopéz-Blanco and Chacón, 2013)*.

### Mouse Experiments

*All experiments were approved by the Institutional Animal Care and Use Committees (IACUC) of and Institutional Biosafety Committee of Yale University (IBSCYU). All the animals were housed under specific pathogen-free conditions in the facilities provided and supported by Yale Animal Resources Center (YARC). hACE2 transgenic B6 mice (heterozygous) were obtained from Jackson Laboratory. 6–8-week-old male and female mice were used for all the experiments. The heterozygous mice were crossed and genotyped to select heterozygous mice for experiments by using the primer sets recommended by Jackson Laboratory*.

### SARS-CoV-2 Infection and Treatment Conditions

*For all in vivo experiments, the 6 to 8 weeks male and female mice were intranasally challenged with 1 x 10^5^ FFU in 25-30 µL volume under anesthesia (0.5 −5 % isoflurane delivered using precision Dräger vaporizer with oxygen flow rate of 1 L/min). For Nab treatment using prophylaxis regimen, mice were treated with 250 µg (12.5 mg/kg body weight) of indicated antibodies (CV3-1 or CV3-25 GASDALIE) via intraperitoneal injection (i.p.) 24 h prior to infection. The starting body weight was set to 100 %. For survival experiments, mice were monitored every 6-12 h starting six days after virus administration. Lethargic and moribund mice or mice that had lost more than 20% of their body weight were sacrificed and considered to have succumbed to infection for Kaplan-Meier survival plots*.

### Focus Forming Assay

*Titers of virus stocks was determined by standard plaque assay. Briefly, the 4 x 10^5^ Vero-E6 cells were seeded on 12-well plate. 24 h later, the cells were infected with 200 µL of serially diluted virus stock. After 1 hour, the cells were overlayed with 1ml of pre-warmed 0.6% Avicel (RC-581 FMC BioPolymer) made in complete RPMI medium. Plaques were resolved at 48 h post infection by fixing in 10 % paraformaldehyde for 15 min followed by staining for 1 hour with 0.2 % crystal violet made in 20 % ethanol. Plates were rinsed in water to visualize plaques*.

### Measurement of Viral Burden

*Indicated organs (nasal cavity, brain, lungs from infected or uninfected mice were collected, weighed, and homogenized in 1 mL of serum free RPMI media containing penicillin-streptomycin and homogenized in 2 mL tube containing 1.5 mm Zirconium beads with BeadBug 6 homogenizer (Benchmark Scientific, TEquipment Inc). Virus titers were measured using three highly correlative methods. Frist, the total RNA was extracted from homogenized tissues using RNeasy plus Mini kit (Qiagen Cat # 74136), reverse transcribed with iScript advanced cDNA kit (Bio-Rad Cat #1725036) followed by a SYBR Green Real-time PCR assay for determining copies of SARS-CoV-2 N gene RNA using primers SARS-CoV-2 N F: 5’-ATGCTGCAATCGTGCTACAA-3’ and SARS-CoV-2 N R: 5’-GACTGCCGCCTCTGCTC-3’*.

*Second, serially diluted clarified tissue homogenates were used to infect Vero-E6 cell culture monolayer. The titers per milligram of tissue were quantified using standard plaque forming assay described above*.

### Analyses of Signature Inflammatory Cytokines mRNA

*Brain and lung samples were collected from mice at the time of necropsy. Approximately, 20 mg of tissue was suspended in 500 µL of RLT lysis buffer, and RNA was extracted using RNeasy plus Mini kit (Qiagen Cat # 74136), reverse transcribed with iScript advanced cDNA kit (Bio-Rad Cat #1725036). To determine levels of signature inflammatory cytokines, multiplex qPCR was conducted using iQ Multiplex Powermix (Bio Rad Cat # 1725848) and PrimePCR Probe Assay mouse primers FAM-GAPDH, HEX-IL6, TEX615-CCL2, Cy5-CXCL10, and Cy5.5-IFNgamma. The reaction plate was analyzed using CFX96 touch real time PCR detection system. Scan mode was set to all channels. The PCR conditions were 95 °C 2 min, 40 cycles of 95 °C for 10 s and 60 °C for 45 s, followed by a melting curve analysis to ensure that each primer pair resulted in amplification of a single PCR product. mRNA levels of Il6, Ccl2, Cxcl10 and Ifng in the cDNA samples of infected mice were normalized to Gapdh with the formula* Δ*C_t_(target gene)=C_t_(target gene)-C_t_(Gapdh). The fold increase was determined using 2^-^*^ΔΔ^*^Ct^ method comparing treated mice to uninfected controls*.

### Virus-Cell Fusion Inhibition Assay

*The split nanoluc assay was used to measure antibody-mediated inhibition of virus-cell fusion (Yamamoto et al., 2019; Lu et al., 2020). Pseudoviruses decorated with SARS-CoV-2 Spike were prepared by transfecting HEK293T cells (70% confluent 10 cm dishes) with a plasmid mixture of 5 µg of psPAX2 (Gag-pol, Rev, and Tat expression vector; does not express Vpr), 5 µg of pCMV-d19 Spike (last 19 residues at C-terminal were deleted) from the B.1.1.7 variant or WH01 G614, and 2 µg of a pCAGGS-Cyclophilin A-HiBiT construct using polyetherimide (PEI). Two days post transfection, virus containing supernatants were clarified using a 0.45 µM PDVF filter (Pall Corp, NY, USA # 4614) and pelleted by ultracentrifugation on a 15% sucrose cushion before resuspension in culture media to achieve a 20X concentration over the original volume. Freshly prepared viruses were incubated for 2 hours at 37°C with triplicate, 10-fold serial dilutions of CV3-25 antibody or non-specific IgG (Jackson Immunoresearch, PA, USA # 305-005-003) in a white, flat bottom 96 well plate (Greiner Bio-One, NC, USA # 655083)*.

*HEK293T-ACE2 target cells were transfected in a 24 well plate using PEI with 500ng/well of pMX Puro PH-LgBiT (LgBiT-tagged to pleckstrin homology domain of human phospholipase Cδ the N terminus). 1 day post transfection, cells were resuspended at 2×10^6^ cells/ml in culture media containing Nano-Glo® Endurazine Live Cell Substrate™ (Promega Inc, WI, USA # N2571) and DrkBiT (Promega Inc, WI, USA # CS3002A01) according to the manufacturer’s recommended concentrations and incubated for 2 hours at 37°C. Labelled target cells were passed through a 70 µM cell strainer and added to the virus + antibody dilution plate (10^5^ cells/well). The assay plate was then incubated for 1 hour at 37°C before measuring luminescence with a Tristar multiwell luminometer (Berthold Technology, Bad Wildbad, Germany). %RLU was calculated by normalizing RLU values to wells without virus (min) and wells without antibody (max)*.

### smFRET Imaging of S on SARS-CoV-2 VLPs

*Lentiviruses carrying SARS-CoV-2 spikes were prepared similarly as previously described (Lu et al., 2020). Two short peptides labeling tags (Q3: GQQQLG; A4: DSLDMLEM) were introduced into designed positions in the S1 subunit on the plasmid encoding S_B.1.1.7_, pCMV-S_B.1.1.7_. Plasmids pCMV-S_B.1.1.7_, dual-tagged pCMV-S_B.1.1.7_ Q3-1 A4-1, and pCMV delta R8.2 were transfected into 293T cells at a ratio of 20:1:21. Using this very diluted ratio of tagged-S vs. wildtype S, for the virus particles containing tagged S, more than 95 % S trimers will have one dual-tagged protomer and two wildtype protomers within a trimer. Using this strategy, we generated lentiviral particles with an average of one dual-tagged S protomer for conjugating FRET-paired fluorophores among predominantly wildtype S trimers presented on lentivirus surface. Viral particles were harvested 40 h post-transfection, filtered with a 0.45 μm pore size filter, and partially purified using ultra-centrifugation at 25,000 rpm for 2 h through a 15 % sucrose cushion made in PBS. Then the particles were re-suspended in 50 mM pH 7.5 HEPES buffer, labeled with self-healing Cy3 and Cy5 derivatives (LD555-CDand LD650-CoA, respectively) and purified through an Optiprep^TM^ (Sigma Aldrich) gradient as previously described (Lu et al., 2019; Lu et al., 2020; Munro et al., 2014). smFRET images of viral particles was acquired on a home-built prism-based total internal reflection fluorescence (TIRF) microscope, as described previously (Lu et al., 2020). The conformational effects of 50 µg/ml CV3-1 and CV3-25 antibodies on SARS-CoV-2 spike were tested by pre-incubating fluorescently labeled viruses for 60 mins at 37°C before imaging in the continued presence of the antibodies. Signals were simultaneously recorded on two synchronized ORCA-Flash4.0 V3 sCMOS cameras (Hamamatsu) at 25 frames per second for 80 seconds. smFRET data analysis was performed using MATLAB (MathWorks)-based customized SPARTAN software package (Juette et al., 2016). Each FRET histogram was fitted into the sum of four Gaussian distributions in Matlab, where each Gaussian distribution represents one conformation and the area under each Gaussian curve estimates the occupancy of each state*.

### Recombinant Protein Expression and Purification

*FreeStyle 293-F (Thermo Fisher) cells were grown to a density of 1×10^6^ cells/mL at 37°C with 8% CO2 with regular 135 rpm agitation. A plasmid encoding for non-cleavable, pre-fusion-stabilized SARS-CoV-2 S ectodomain (1-1208) (HexaPro, S-6P (Hsieh et al., 2020; Wrapp et al., 2020) - a gift from Dr. Jason S. McLellan) with a removable C-terminal twin-strep tag was transfected into cells with EndoFectin Max (GeneCopoeia) using the manufacturer’s protocol. One-week post-transfection, the clarified supernatant was purified on strep-tactin resin (IBA) followed by size-exclusion chromatography on a Superose 6 10/300 column (GE Healthcare) equilibrated with 10 mM Tris-HCl pH 8.0 and 200 mM NaCl as the running buffer (SEC buffer). The C-terminal twin-Strep-Tag was removed by HRV3C (Sigma Aldrich) digestion overnight at 4 °C and the uncleaved protein was removed by passage over Ni-NTA resin. The cleaved protein was further purified on a Superose 6 10/300 column in SEC buffer. Alternatively, cells were transfected with a plasmid coding for SARS-CoV-2 RBD or ACE2-Fc and were purified on Ni-NTA resin (Invitrogen) or Protein A resin (Cytiva), respectively. Protein purity was confirmed by SDS-PAGE. Only freshly isolated protein was used for Cryo-EM grid preparations*.

*Expression plasmids encoding the heavy and light chains of CV3-1 IgG or CV3-25 IgG were transiently transfected into Expi293F cells (Thermo Fisher) with ExpiFectamine 293 transfection reagent using the manufacturer’s protocol (Thermo Fisher). After 6-days post transfection, antibody was purified on Protein A resin from cell supernatant (Thermo Fisher). Fab was generated by overnight papain digestion at 37°C using immobilized papain agarose (Thermo Fisher). Fab was separated from Fc and uncleaved IgG by passage over protein A resin followed by size-exclusion chromatography on a Superose 6 10/300 column before being used in SPR binding, X-Ray crystallography or Cryo-EM experiments*.

### Surface Plasmon Resonance

*All surface plasma resonance assays were performed on a Biacore 3000 (GE Healthcare) with a running buffer of 10 mM HEPES pH 7.5 and 150 mM NaCl supplemented with 0.05% Tween 20 at 25°C. Initial peptide scanning was performed by the binding of a series of SARS-CoV-2 S2 synthetic peptides (GenScript) to immobilized CV3-25 IgG (∼5800 RU) on a Protein A sensor chip (Cytiva). For the kinetic binding measurements of S2 peptides #289 (15-mer), #289 (11-mer) and the 26mer (1140-1165) to CV3-25, ∼5800 RU of CV3-25 IgG was first immobilized on a protein A chip (Cytiva) and 2-fold serial dilutions of the S2 peptides were then injected with concentrations ranging from 6.25 to 200 nM. After each cycle the protein A sensor chip was regenerated with 0.1 M Glycine pH 2.0. CV3-1 IgG was used as a negative control. All sensorgrams were corrected by subtraction of the corresponding blank channel in addition to the buffer background and the kinetic constant determined using a 1:1 Langmuir model with the BIAevaluation software (GE Healthcare). Goodness of fit of the curve was evaluated by the Chi2 value with a value below 3 considered acceptable*.

### Cryo-EM Sample Preparation and Data Collection

*The purified non-tagged SARS-CoV-2 HexaPro spike (293F produced) was incubated with 20-fold excess of CV3-25 Fab overnight at 4°C before purification on a Superose 6 300/10 GL column (GE Healthcare). The complex peak was harvested, concentrated to about 0.5 mg/mL in SEC buffer and immediately used for CryoEM grid preparation. 3μL of protein was deposited on a holey copper grids (QUANTIFOIL R 1.2/1.3, 200 mesh,* EMS) *which had been glow-discharged for 30s at 15 mA (Tedpella Inc). The grids were vitrified in liquid ethane using a Vitrobot Mark IV (Thermo Fisher) with a blot time of 2-4 s and the blot force of 20 at 4 °C and 95% humidity*.

*Cryo-EM data from a good grid were acquired in 300kV Titan Krios electron microscope, equipped with a Gatan K2-BioQuantum Image filter camera system (Thermo Fisher and Gatan Inc.) in National Cancer Institute/NIH IRP cryoEM facility, Bethesda MD. 50-frame image stacks were collected at a magnification of 165,000x, corresponding to a calibrated pixel size of 0.821 Å/pixel, with a total exposure dose of 59.3 e^-^/ Å from 5s exposure*.

### CryoEM Data Processing, Model building and Analysis

*Motion correction, CTF estimation, particle picking, curation and extraction, 2D classification, ab initio model reconstruction, volume refinements and local resolution estimation were carried out in cryoSPARC (Punjani et al., 2017; Rubinstein and Brubaker, 2015). An initial SARS-CoV-2 spike model (PDB: 6XKL (Hsieh et al., 2020)) with single-RBD up was used as a modeling template. The NTDs were initially modelled from PDB entry 7LY3(McCallum et al., 2021). The initial docking model for CV3-25 Fab was taken from the crystallography model in this study*.

*Automated and manual model refinements were iteratively carried out in ccpEM (Burnley et al., 2017), Phenix (real-space refinement) (Liebschner et al., 2019) and Coot(Emsley and Cowtan, 2004). Geometry validation and structure quality evaluation were performed by EM-Ringer (Barad et al., 2015) and Molprobity (Chen et al., 2010). Model-to-map fitting cross correlation and figures generation were carried out in USCF Chimera, Chimera X (Goddard et al., 2018; Pettersen et al., 2004; Pettersen et al., 2021) and PyMOL (The PyMOL Molecular Graphics System, Version 2.0 Schrödinger, LLC.). The complete cryoEM data processing workflow is shown in* Figure S2 *and statistics of data collection, reconstruction and refinement is described in* Table S2.

### Crystallization and Structure Determination of CV3-25 with S2 Stem Peptide

*CV3-25 Fab was prepared and purified as described(Ullah et al., 2021). 10 mg/mL of CV3-25 was mixed with synthetic S2 peptide spanning residues 1153-1163, 1153-1167 or 1140-1165 (26mer) in a 1:10 molar ratio of Fab to peptide. Crystal screening of Fab-peptide complexes were performed using the vapor-diffusion hanging drop method using the sparse matrix crystallization screens ProPlex (Molecular Dimensions), Index (Hampton Research), or Crystal Screen I and II (Hampton Research) with a 1:1 ratio of protein to well solution. After approximately 2 weeks incubation at 21 °C, diffraction-quality co-crystals of the Fab-26mer were obtained in 0.1 M sodium citrate pH 5.6, 20% PEG4000 and 20% isopropanol. Crystals were snap-frozen in the crystallization condition supplemented with 20% 2-methyl-2, 4-pentanediol (MPD) as the cryoprotectant. X-ray diffraction data was collected at the SSRL beamline 9-2 and was processed with HKL3000(Minor et al., 2006). The structure was solved by molecular replacement in Phenix(Liebschner et al., 2019) using a CV3-25 framework model generated by SAbPred(Dunbar et al., 2016). Iterative cycles of model building and refinement were done in Coot(Emsley and Cowtan, 2004) and Phenix. Structural analysis and figure generation were performed in PyMOL and ChimeraXFab-peptide interface and buried surface area were determined in PISA(Krissinel and Henrick, 2007). Data collection and refinement statistics are shown in Table 1*.

### Flow Cytometry Analysis of Cell-Surface Staining

*Using the standard calcium phosphate method, 10* μg *of Spike expressor and 2* μg *of a green fluorescent protein (GFP) expressor (pIRES2-eGFP; Clontech) was transfected into 2 × 10^6^ 293T cells. At 48h post transfection, 293T cells were stained with anti-Spike monoclonal antibodies CV3-25, CV3-1 (5 µg/mL) or using the ACE2-Fc chimeric protein (20 µg/mL) for 45 min at 37°C. Alternatively, to determine the Hill coefficients (Anand et al., 2020), cells were preincubated with increasing concentrations of CV3-25 or CV3-1 (0.04 to 20 µg/mL). Alexa Fluor-647-conjugated goat anti-human IgG (H+L) Abs (Invitrogen) were used as secondary antibodies to stain cells for 30 min at room temperature. The percentage of transfected cells (GFP+ cells) was determined by gating the living cell population based on the basis of viability dye staining (Aqua Vivid, Invitrogen). Samples were acquired on a LSRII cytometer (BD Biosciences) and data analysis was performed using FlowJo v10.5.3 (Tree Star). Hill coefficient analyses were done using GraphPad Prism version 9.1.0 (GraphPad). Alternatively, for peptide epitope competition assay, CV3-25 (5µg/mL) was pre-incubated in presence of increasing concentrations of peptide #288 (1149-KEELDKYFKNHTSPD-1163), peptide #289 (1153-DKYFKNHTSPDVDLG-1167), a shorter version of peptide #289 (1153-DKYFKNHTSPD-1163) or a scramble version of the peptide #289 (DHDTKFLNYDPVGKS), which were synthesized by Genscript*.

### Viral Neutralization Assay

*293T-ACE2 target cells were infected with single-round luciferase-expressing lentiviral particles (Prévost et al., 2020). Briefly, 293T cells were transfected by the calcium phosphate method with the lentiviral vector pNL4.3 R-E-Luc (NIH AIDS Reagent Program) and a plasmid encoding for SARS-CoV-2 Spike at a ratio of 5:4. Two days post-transfection, cell supernatants were harvested and stored at –80°C until further use. 293T-ACE2 target cells were seeded at a density of 1×10^4^ cells/well in 96-well luminometer-compatible tissue culture plates (Perkin Elmer) 24h before infection. To measure virus neutralization, recombinant viruses in a final volume of 100 μL were incubated with increasing concentrations of CV3-1 or CV3-25 (0.01 to 10 µg/mL) for 1h at 37°C and were then added to the target cells followed by incubation for 48h at 37°C; cells were lysed by the addition of 30 μL of passive lysis buffer (Promega) followed by one freeze-thaw cycle. An LB942 TriStar luminometer (Berthold Technologies) was used to measure the luciferase activity of each well after the addition of 100 μL of luciferin buffer (15 mM MgSO_4_, 15 mM KH_2_PO_4_ [pH 7.8], 1 mM ATP, and 1 mM dithiothreitol) and 50 μL of 1 mM D-luciferin potassium salt (Prolume). The neutralization half-maximal inhibitory dilution (IC_50_) represents the antibody concentration inhibiting 50% of the infection of 293T-ACE2 cells by recombinant viruses bearing the indicated surface glycoproteins. Alternatively, for peptide epitope competition assay, CV3-25 (10µg/mL) was pre-incubated in presence of increasing concentrations of peptide #289 (1153-DKYFKNHTSPDVDLG-1167) or a scramble version of the same peptide (DHDTKFLNYDPVGKS)*.

### Radioactive Labeling and Immunoprecipitation

*For pulse-labeling experiments, 5 × 10^5^ 293T cells were transfected by the calcium phosphate method with SARS-CoV-2 Spike expressors. One day after transfection, cells were metabolically labeled for 16 h with 100μCi/ml [^35^S]methionine-cysteine ([^35^S] protein labeling mix; Perkin-Elmer) in Dulbecco’s modified Eagle’s medium lacking methionine and cysteine and supplemented with 10% of dialyzed fetal bovine serum and 1X GlutaMAX^TM^ (ThermoFisher Scientific). Cells were subsequently lysed in radioimmunoprecipitation assay (RIPA) buffer (140 mM NaCl, 8 mM Na_2_HPO_4_, 2 mM NaH_2_PO_4_, 1% NP-40, 0.05% sodium dodecyl sulfate [SDS], 1.2mM sodium deoxycholate [DOC]) with protease inhibitors (ThermoFisher Scientific). Precipitation of radiolabeled SARS-CoV-2 Spike glycoproteins from cell lysates or supernatant was performed with CV3-25 in combination with a polyclonal rabbit antiserum raised against SARS-CoV-2 RBD protein for 1 h at 4°C in the presence of 45 μL of 10% protein A-Sepharose beads (GE Healthcare)*.

### Peptide Scanning ELISA

*SARS-CoV-2 Spike peptide ELISA (enzyme-linked immunosorbent assay) The SARS-CoV-2 Spike ELISA assay used was adapted from a previously described ELISA (Prevost et al., 2020). Peptides covering the entire SARS-CoV-2 S2 sequence with a length of 15 residues (15-mer) and an overhang of 4 residues were purchased from JPT Peptide Technologies. Briefly, SARS-CoV-2 S2 peptide pools or individual peptides (1 μg/ml), or bovine serum albumin (BSA) (1μg/ml) as a negative control, were prepared in PBS and were adsorbed to plates (MaxiSorp; Nunc) overnight at 4 °C. Coated wells were subsequently blocked with blocking buffer (Tris-buffered saline [TBS] containing 0.1% Tween20 and 2% BSA) for 1 hour at room temperature. Wells were then washed four times with washing buffer (TBS containing 0.1% Tween20). CV3-25 mAb (50 ng/ml) was prepared in a diluted solution of blocking buffer (0.1 % BSA) and incubated with the peptide-coated wells for 90 minutes at room temperature. Plates were washed four times with washing buffer followed by incubation with HRP-conjugated anti-IgG secondary Abs (Invitrogen) (diluted in a diluted solution of blocking buffer [0.4% BSA]) for 1 hour at room temperature, followed by four washes. HRP enzyme activity was determined after the addition of a 1:1 mix of Western Lightning oxidizing and luminol reagents (Perkin Elmer Life Sciences). Light emission was measured with a LB942 TriStar luminomete (Berthold Technologies). Signal obtained with BSA was subtracted for each plate*.

### Western Blotting

*293T-S cells express the wild-type S glycoprotein from a SARS-CoV-2 Wuhan-Hu-1 strain (Nguyen et al., 2021). 293T-S cells were seeded in 6-well plates at a density of 1×10^6^ cells per well on day 0. On day 1, cells were either induced with 1 µg/ml doxycycline or mock treated as a control. Two days after induction, cells were lysed with lysis buffer (1x PBS, 1% NP-40, 1x protease inhibitor cocktail (Roche)). Cell lysates were subjected to Western blotting using the CV3-1 or CV3-25 antibodies; mouse anti-S1 antibody (Sino Biological) and rabbit anti-S2 antibody (Sino Biological) were used as controls. The Western blots were developed with horseradish peroxidase (HRP)-conjugated secondary antibodies (anti-human IgG, anti-mouse IgG or anti-rabbit IgG, correspondingly). To evaluate antibody recognition of S glycoproteins lacking N-linked glycans, 293T-S cells expressing the wild-type SARS-CoV-2 S glycoprotein were lysed with lysis buffer, as described above. Lysates were treated with PNGase F (NEB) following the manufacturer’s instructions or mock treated as a control. The lysates were then Western blotted with the CV3-25 antibody, as described above*.

### Quantification and Statistical Analysis

*Data were analyzed and plotted using GraphPad Prism software (La Jolla, CA, USA). Statistical significance for pairwise comparisons were derived by applying non-parametric Mann-Whitney test (two-tailed). To obtain statistical significance for survival curves, grouped data were compared by log-rank (Mantel-Cox) test. To obtain statistical significance for grouped data we employed 2-way ANOVA followed by Tukey’s multiple comparison tests*.

*p values lower than 0.05 were considered statistically significant. P values were indicated as *, p < 0.05; **, p < 0.01; ***, p < 0.001; ****, p < 0.0001*.

### Schematics

*Schematics for showing experimental design in figures were created with BioRender.com*.

